# CXCL4 links inflammation and fibrosis through transcriptional and epigenetic reprogramming of monocyte-derived cells

**DOI:** 10.1101/807230

**Authors:** Sandra C Silva-Cardoso, Weiyang Tao, Chiara Angiolilli, Ana P. Lopes, Cornelis P.J. Bekker, Abhinandan Devaprasad, Barbara Giovannone, Jaap van Laar, Marta Cossu, Wioleta Marut, Erik Hack, Rob J de Boer, Marianne Boes, Timothy R.D.J. Radstake, Aridaman Pandit

**Affiliations:** Center for Translational Immunology, Department of Immunology, University Medical Center Utrecht, Utrecht University, The Netherlands; Department of Rheumatology and Clinical Immunology, University Medical Center Utrecht, Utrecht University, The Netherlands; Department of Dermatology and Allergology, University Medical Center Utrecht, Utrecht University, The Netherlands; Theoretical Biology, University Utrecht, The Netherlands; Department of Pediatrics, University Medical Center Utrecht, Utrecht University, The Netherlands

## Abstract

Fibrosis is a condition shared by numerous inflammatory diseases. Our incomplete understanding of the molecular mechanisms underlying fibrosis has severely hampered effective drug development. CXCL4 is associated with the onset and extent of fibrosis development in systemic sclerosis, a prototypic inflammatory and fibrotic disease. Here, we integrated 65 paired longitudinal transcriptional and methylation profiles from monocyte-derived cells with/without CXCL4 exposure. Using data-driven gene regulatory network analyses, we demonstrate that CXCL4 dramatically alters the trajectory of monocyte differentiation, inducing a novel pro-inflammatory and pro-fibrotic phenotype mediated via key transcriptional regulators including CIITA. CXCL4 exposed monocyte-derived cells directly trigger a fibrotic cascade by producing ECM molecules and inducing myofibroblast differentiation. Inhibition of CIITA mimicked CXCL4 in inducing a pro-inflammatory and pro-fibrotic phenotype, validating the relevance of the gene regulatory network. Our study unveils CXCL4 as a key secreted factor driving innate immune training and forming the long-sought link between inflammation and fibrosis.

## Introduction

Fibrosis is uncontrolled accumulation of extracellular matrix (ECM) in multiple organs and accounts for one third of deaths worldwide(Rinkevich et al., 2015; Zeisberg & Kalluri, 2013). Fibrosis is considered to be a result of complex cellular and molecular interplay following tissue inflammation and injury. Across a wide range of diseases, fibroblasts inappropriately synthesize and release increased amounts of ECM components, suggesting a conceptual framework in which myofibroblast transition is the key event leading to fibrosis(Rinkevich et al., 2015). Recent studies however, strongly implicate the innate immune system as a critical contributor to fibrosis development(Wick et al., 2010), in line with clinical observations that an inflammatory phase precedes fibrosis by years. Hence, identification of the molecular pathways linking inflammation to fibrosis will provide unprecedented opportunities for drug development to treat or even reverse tissue fibrogenesis(Wick et al., 2010; Zeisberg & Kalluri, 2013).

CXCL4, a chemokine initially identified as a product of activated platelets, is now known to be secreted by a variety of immune cells(Levine & Wohl, 1976; Schaffner, Rhyn, Schoedon, & Schaer, 2005; van Bon, Affandi, et al., 2014). CXCL4 drives a broad spectrum of immune-modulatory effects in both hematopoietic stem and progenitor cells, as well as differentiated immune cells, and has been implicated in the pathology of a variety of inflammatory diseases(Affandi et al., 2018; Kioon et al., 2018; Scheuerer et al., 2000; Silva-Cardoso et al., 2017; Silva-Cardoso, Bekker, Boes, Radstake, & Angiolilli, 2019). Systemic sclerosis (SSc) is an archetypical fibrotic disease in which hypoxia, followed by endothelial cell damage and immune activation, typically culminates in fibrosis of the skin and internal organs. Previously, we identified CXCL4 as an early biomarker of this process(van Bon, Affandi, et al., 2014). Monocytes are indispensable for inflammation and repair. Monocyte-derived dendritic cells (moDCs) can be differentiated *in vitro* by culturing monocytes isolated from human donors and are considered as DC model that mimics *in vivo* DC biology. Previously, we investigated whether circulating CXCL4 potentiates aberrant TLR-mediated responses and T-cell dysregulated responses observed in autoimmune diseases including SSc (Affandi et al., 2018; Silva-Cardoso et al., 2017). Considering the presence of CXCL4 during early inflammation and its role in modulating key immune functions, we postulated that CXCL4 might constitute the link between inflammation and fibrosis. Therefore, here, we tested this hypothesis by examining the transcriptional and epigenetic effects of CXCL4 on monocytes during and after differentiation, integrating 65 paired time courses of whole genome transcriptional and methylation profiles, and reconstructed CXCL4-dependent gene regulatory networks.

## Results

### CXCL4 drastically impacts monocyte differentiation

To study the role of CXCL4 on the possible imprinting of immune cells towards fibrosis, we examined the effects of CXCL4 on the differentiation of monocyte-derived dendritic cells (moDCs)(Silva-Cardoso et al., 2017). We cultured monocytes obtained from five healthy donors in the presence of IL-4 and GM-CSF to differentiate them in the absence (conventional moDCs) and presence of CXCL4 (“CXCL4-moDCs”). To systematically study the effects of CXCL4 on the trajectory of monocyte differentiation into moDCs, we obtained longitudinal transcriptional (RNA-seq) profiles at days 0 (monocytes), 2, 4, and 6. To examine the effects of CXCL4 on moDC maturation, we stimulated the cells on day 7 with the toll-like receptor 3 ligand polyI:C and obtained transcriptional profiles before stimulation (day 7), 4 hours (day 7 + 4 hours) and 24 hours (day 8) after stimulation (**Figure 1A**).

**Figure 1.**
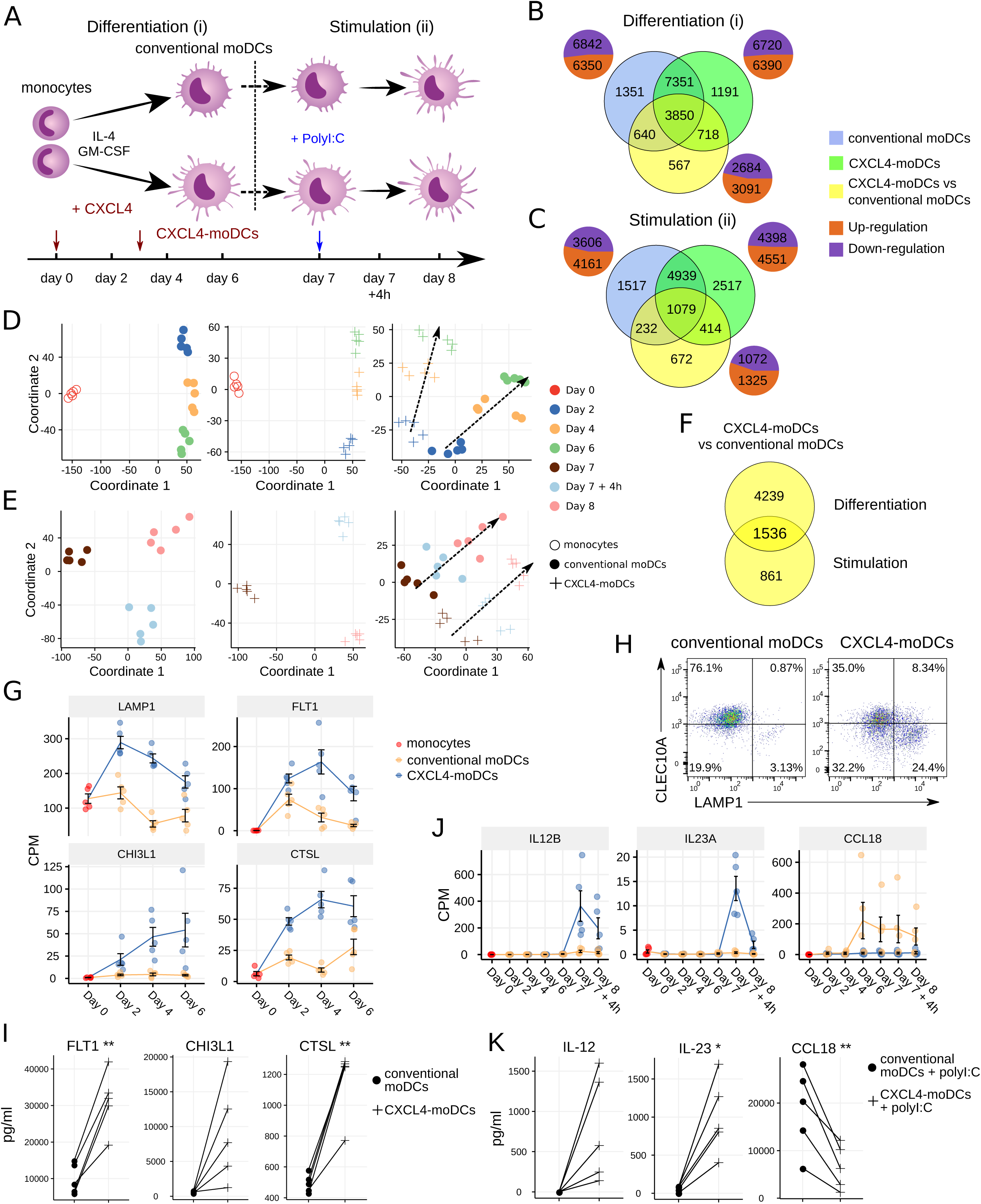
Transcriptomic programing of CXCL4-moDCs. **(A)** Schematic overview of the experimental setup: (i) differentiation of monocytes to conventional moDCs or CXCL4-moDCs; (ii) stimulation with polyI:C on day 7, for 4 hours or 24 hours. Overlap of differentially expressed genes (DEGs) during **(B)** differentiation and **(C)** after polyI:C stimulation of: monocytes into conventional moDCs (blue); monocytes into CXCL4-moDCs (green); and between CXCL4-moDCs and moDCs during differentiation (yellow). In **(B)** and **(C)** pie charts showing the number of upregulated (orange) and down-regulated (purple) genes. Multi-dimensional scaling (MDS) plot **(D)** differentiating and **(E)** stimulated conventional moDCs (left panel), CXCL4-moDCs (middle panel), and CXCL4-moDCs vs conventional moDCs (right panel). In **(D)** and **(E)** dotted lines indicate trajectories over time. **(F)** Overlap of DEGs between CXCL4-moDCs and conventional moDCs, during differentiation and upon stimulation. Gene expression of example genes differential during **(G)** differentiation and **(J)** stimulation between CXCL4-moDCs and conventional moDCs. Validation of **(H)** protein expression (flow cytometry) and **(I)** cytokine production (Luminex) on day 6. **(K)** Validation of cytokine production (Luminex) on day 8. Gene expressions are shown as mean±SEM. CPM, count per million. In panels I and K, lines connect individual donors (n=5). \**P*<0.05; \*\**P*< 0.01, paired two-sided Student’s *t*-test.

The differentiation of moDCs from monocytes was accompanied by extensive transcriptional changes, as 13,192 genes underwent significant (likelihood ratio test; FDR corrected p-value ≤ 0.05) alterations in their expression levels (**Figure 1B**). Nearly half of the differentially expressed genes (6,350) were upregulated. CXCL4-moDCs also underwent widespread transcriptional changes, as 13,110 genes were differentially expressed compared to monocytes, nearly half of those were found to be upregulated (**Figure 1B** and **Figure 1—figure supplement 1**). Remarkably, most of this transcriptional shift happens between day 0 and day 2, in both conventional moDCs and CXCL4-moDCs (**Figure 1D**). Genes characteristic of monocyte differentiation (such as *CD14*, *CD163*, *TLR2*, *TLR4*, and *TLR7*)(Schinnerling, Garcia-Gonzalez, & Aguillon, 2015) and cell adhesion molecules (including *LGALS2*, *LGALS9*, and *ICAM2*) were down-regulated in both conventional moDCs and CXCL4-moDCs on day 2 (**Supplementary File 3**). After day 2, cells continued to differentiate, as evidenced by their shifting transcriptional profiles (**Figure 1D**). Genes encoding pattern recognition proteins MRC1, MRC2, growth factors such as CSF1, and the chemokine receptors CCR1, CCR5, and CCR7, were upregulated over time in both conventional moDCs and CXCL4-moDCs (**Supplementary File 3**). Together these results indicate that the differentiation of monocytes (with or without CXCL4) leads to massive transcriptional changes as reported by several previous studies(Gleissner, Shaked, Little, & Ley, 2010; Vento-Tormo et al., 2016).

To elicit the transcriptional signature unique to CXCL4 exposure, we compared differentiating CXCL4-moDCs with conventional moDCs from day 2 to day 6 and found differential expression of 5,775 genes (likelihood ratio test, FDR corrected p-value ≤ 0.05; **Figure 1B, Figure 1—figure supplement 1**, and **Supplementary File 1 and 3**). CXCL4-moDCs follow a distinct molecular differentiation trajectory that progressively diverges from conventional moDCs (**Figure 1D**, right panel). The CXCL4 signature genes belong to several crucial innate immune system pathways including cytokine signaling, interferon signaling, and antigen processing and presentation (**Figure 1—figure supplement 1E**). For instance, CXCL4-moDCs, in the absence of further stimulation, up-regulate expression of several inflammatory molecules such as *CTSL*, *FLT1*, *CD86*, *LAMP1*, *CHI3L1*, and down-regulate signaling receptors such as *CLEC10A, IL1R1, IL1R2* compared to moDCs (**Figure 1G-I** and **Figure 1—figure supplement 1A-D**). Strikingly, CXCL4 exposure also leads to dramatic changes in expression of genes regulating metabolism and transcription (**Figure 1—figure supplement 1E**), reminiscent of changes previously observed in myeloid cells undergoing immune training(Saeed et al., 2014). Thus, CXCL4 orchestrates a differentiation process dramatically different than that of the conventional moDCs.

### Mature CXCL4-moDCs are functionally distinct from conventional moDCs

To study the effects of CXCL4 on moDC maturation, we stimulated the cells with polyI:C on day 7. This perturbed the expression of 8,949 and 7,767 genes in CXCL4-moDCs and conventional moDCs, respectively, compared to the day 7 transcriptional profiles of their unstimulated counterparts (**Figure 1C** and **1E**, left and middle panels). 2,397 genes responded differently to polyI:C stimulation in CXCL4-moDCs compared to conventional moDCs (**Figure 1C** and **Figure 1—figure supplement 2**). Several pathways involved in inflammatory responses such as TLR signaling, interferon signaling, and cytokine signaling, were significantly upregulated in CXCL4-moDCs compared to conventional moDCs (**Figure 1—figure supplement 2E**). Confirming our previous findings(Silva-Cardoso et al., 2017), these transcriptional changes were followed by increased production of pro-inflammatory mediators such as IL-1β, IL-6, IL-12, IL-23, IL-27, TNF and CCL22, and down-regulation of immune-suppressive mediator CCL18 (validated using Luminex assays; see **Figure 1J-K** and **Figure 1— figure supplement 2A-D**). Pathways involved in cellular adhesion, integrin signaling, ECM organization, and collagen formation, among others, were upregulated in CXCL4-moDCs upon polyI:C stimulation compared to stimulated moDCs (**Figure 1—figure supplement 2E**), indicating that CXCL4 exposure induces a pro-inflammatory and pro-fibrotic phenotype. Because most of the altered genes were already differentially expressed in immature CXCL4-moDCs (**Figure 1E, F** and **Figure 1—figure supplement 2B**), the unique molecular program induced by CXCL4 is suggestive of genetic imprinting.

### CXCL4 alters epigenetic imprinting during differentiation but not maturation of moDCs

To comprehensively examine whether CXCL4 signaling might alter moDC phenotype via epigenetic modifications(Broen, Radstake, & Rossato, 2014; Vento-Tormo et al., 2016; Zhang et al., 2014), we studied genome wide alterations in DNA methylation. Similar to the transcriptome analysis, we found that a large number of genes, regions and sites were differentially methylated between monocytes and differentiating moDCs and CXCL4-moDCs (**Figure 2A**, **Figure 2—figure supplement 1**). Interestingly, most of the differentially methylated genes were hypomethylated compared to monocytes (2,617 in conventional moDCs and 2,156 in CXCL4-moDCs) (**Figure 2A-B**, **Figure 2—figure supplement 1A and C**, and **Supplementary File 4**).

**Figure 2.**
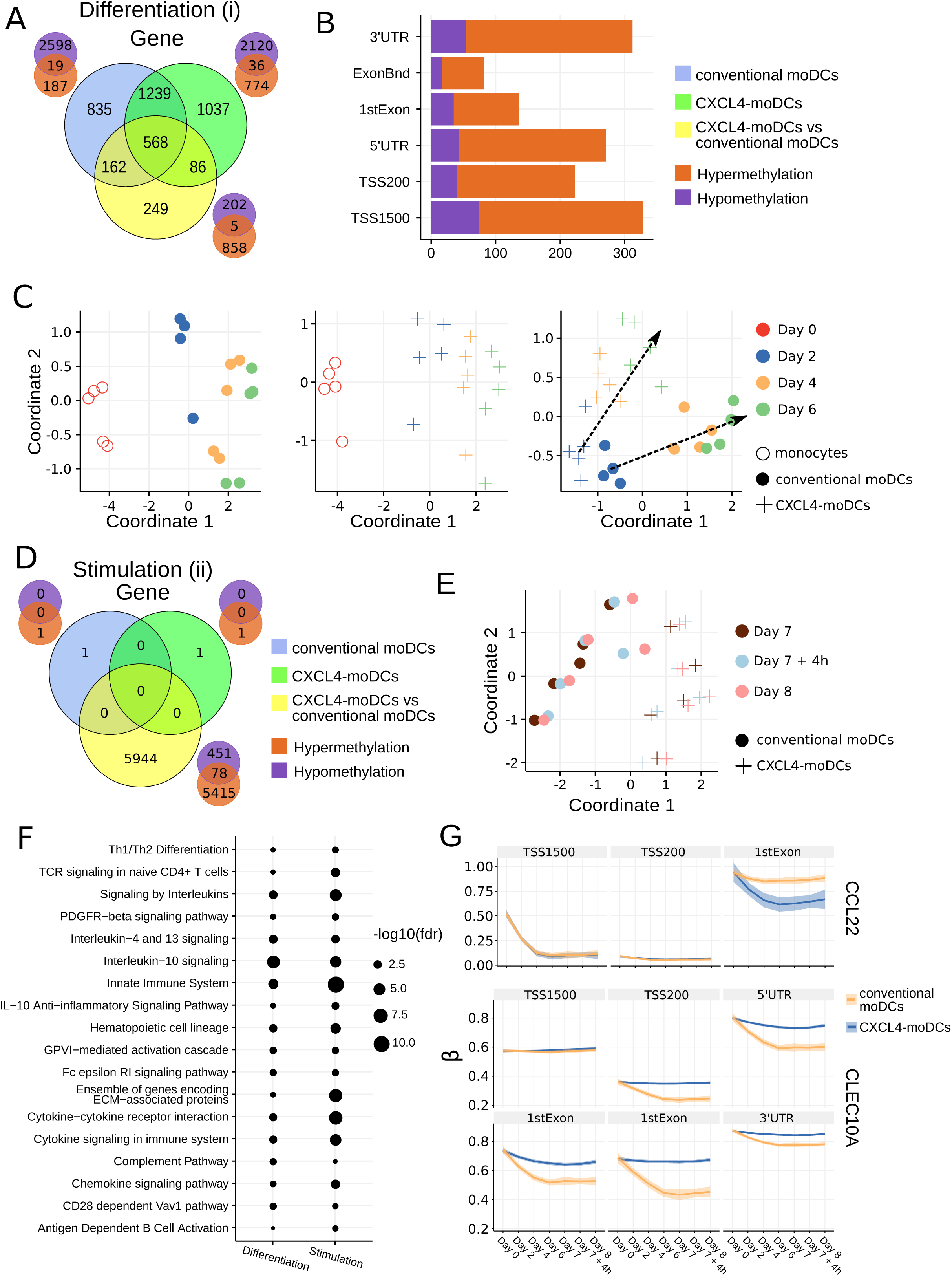
DNA methylation analysis of CXCL4-moDCs and conventional moDCs. **(A)** Overlap between differentially methylated genes (DMGs) found during differentiation similar to Figure 1B. A gene is considered differentially methylated if any region on the gene is differentially methylated. Smaller Venn diagram graphs display the overlap of hyper-methylated (orange) and hypo-methylated (purple) genes for each comparison. Note some genes are classified as both hyper-methylated and hypo-methylated based on different regions. **(B)** Distribution of differentially methylated regions (1500 and 200 base pairs upstream of the transcription start site (TSS), 5’ untranslated region (UTR), 1st exon, other exons (ExonBnd) and 3’ UTR) between CXCL4-moDCs and conventional moDCs during differentiation. **(C)** MDS analysis using DMRs, similar to Figure 1D. **(D)** Overlap between DMGs found during stimulation similar to Figure 1C. **(E)** MDS analysis using all DMRs between CXCL4-moDCs and conventional moDCs during stimulation. **(F)** Top enriched pathways from DMGs between CXCL4-moDCs and conventional moDCs during differentiation and stimulation. **(G)** DNA methylation β values (see Methods) of CCL22 and CLEC10A. Lines represent mean β values and shading represents 95% confidence interval.

To discern the epigenetic footprint of CXCL4 during differentiation, we compared the methylome profiles of differentiating CXCL4-moDCs with differentiating conventional moDCs (from day 2 to day 6). CXCL4 exposure led to substantial changes in the DC methylome as 1,065 genes were differentially methylated between CXCL4-moDCs and conventional moDCs (**Figure 2A** and **Figure 2—figure supplement 1**). Most of the differentially methylated genes were hypermethylated in CXCL4-moDCs. The hypermethylation was not restricted to promoter regions, indicating that CXCL4 influences chromatin accessibility at a more global level (**Figure 2B**). Alterations in DNA methyltransferases and DNA demethylases are known to cause global hypermethylation, which have been implicated in SSc pathogenesis previously(Altorok, Tsou, Coit, Khanna, & Sawalha, 2015). Interestingly, we found transcriptional upregulation of DNA methyltransferases (such as DNMT3A) and downregulation of DNA demethylases (TET2 and TET3) that together can cause global hypermethylation in CXCL4-moDCs (**Figure 3—figure supplement 1A**). As in the transcriptional analysis, CXCL4-moDCs progressively diverge from moDCs (**Figure 2C**, right panel). This progressive and temporal divergence of DNA methylation patterns caused by CXCL4 alters several crucial innate immune system pathways including cytokine signaling, co-stimulatory molecules, and ECM organization (**Figure 2F**). Thus, we provide direct mechanistic evidence that CXCL4 programs a pro-inflammatory and pro-fibrotic phenotype via epigenetic imprinting that corroborates the transcriptional results (**Figure 2—figure supplement 1E-F**).

We also studied the role of epigenetic remodeling in mature moDCs. Surprisingly, stimulation of conventional moDCs and CXCL4-moDCs with polyI:C on day 7 hardly affected the DNA methylation (**Figure 2D, G**, **Figure 2—figure supplement 1B and D**), an observation confirmed by multivariate analysis as the samples did not exhibit any temporal clustering (**Figure 2E**). Thus, the altered functional responses exhibited by CXCL4-moDCs were epigenetically imprinted during differentiation rather than maturation (**Figure 2—figure supplement 1**).

### Gene regulatory network driving the CXCL4-specific transcriptome

Since CXCL4 exposure caused massive alterations in both DNA methylation and transcriptional factors, we next studied the regulatory mechanisms behind the CXCL4 signature. We first assessed the concordance of DNA methylation and mRNA expression and found that the changes in DNA methylation did not correlate with the changes in corresponding gene’s expression for majority of the CXCL4 signature genes (**Figure 3A** and **Figure 3—figure supplement 1B-E**). Surprised by the lack of correlations for most CXCL4 signature genes, we checked if levels of DNA methylation reflect upon the overall gene expression levels, rather than their differential expression. We indeed found that levels of DNA methylation play a role in overall gene expression levels (**Figure 3—figure supplement 1F-G**). Given the lack of concordance between individual transcriptional and DNA methylation changes, DNA methylation of individual genes may be a poor guide for further association studies.

**Figure 3.**
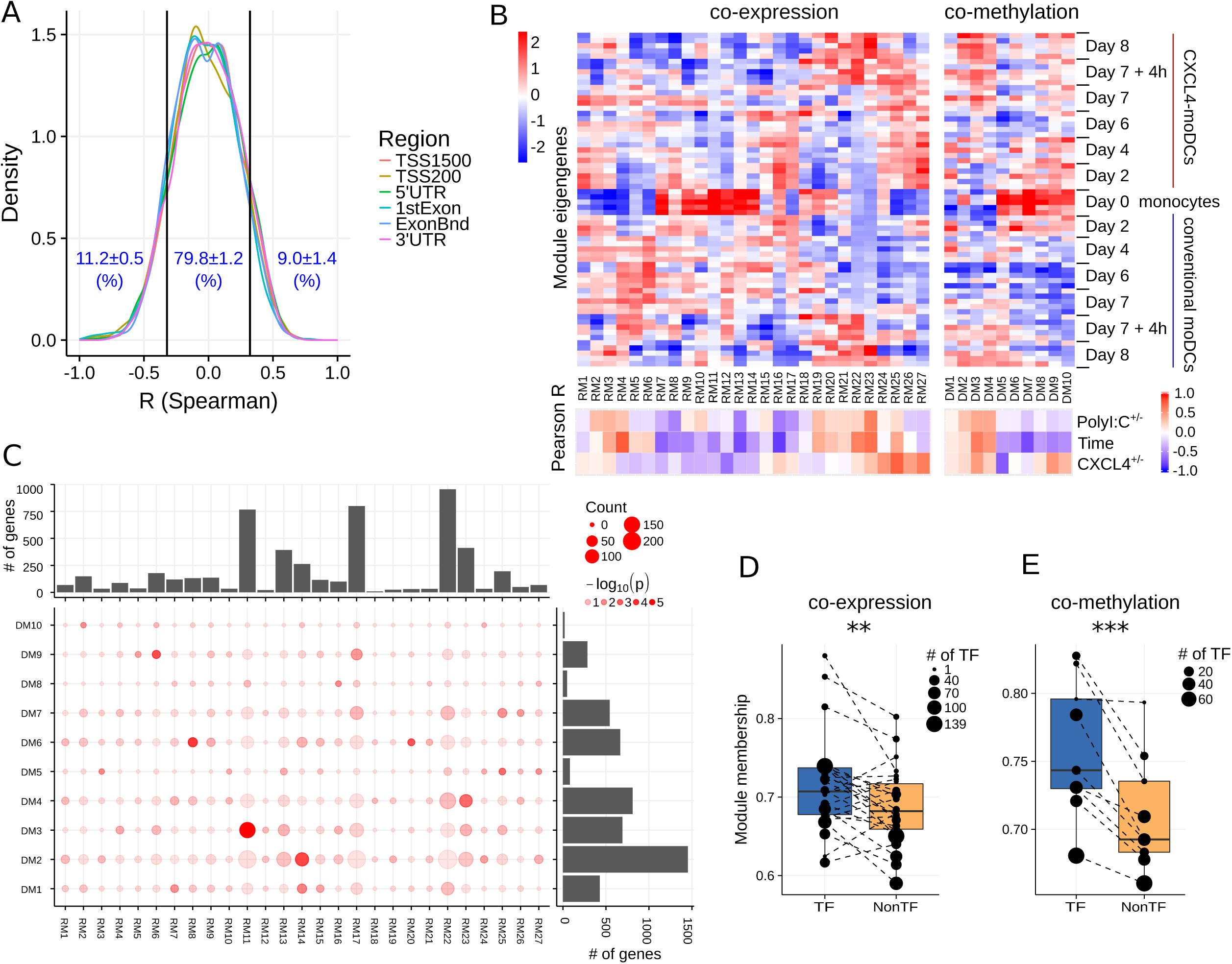
Co-expression and co-methylation networks. **(A)** Distribution of spearman correlation coefficients (R) between β values of each region and the corresponding gene expression for all genes that are differentially expressed and methylated. The cutoffs (two vertical lines at R = ±0.32) indicate significant correlation coefficients (p < 0.01). **(B)** The top heatmap shows expression/methylation eigengenes of co-expression (left) and co-methylation (right) modules. The bottom heatmap shows the Pearson correlation coefficients between sample traits (i.e. CXCL4^+/-^, time and polyI:C^+/-^), and co-expression (left) and co-methylation (right) module eigengenes. **(C)** Concordance of co-expression and co-methylation modules. The bottom left graph shows the number (circle size) and significance (color, p-value calculated by Fisher’s exact test) of overlapping genes between co-expression and co-methylation modules. The bar plots show the total number of genes in the co-expression (top) or co-methylation (right) module. Module membership comparisons of transcriptional regulators (TF) and other genes (NonTF) in **(D)** co-expression and **(E)** co-methylation network. Each dot represents a module and the size denotes the number of TFs in the corresponding module. Modules that do not contain TFs were excluded in these analyses. \*\**P*<0.01, \*\*\**P*<0.001, paired two-sided Student’s *t*-test.

Genes rarely work in isolation, and their expression is typically regulated via a complex molecular network(Goode et al., 2016; Ramirez et al., 2017). To systematically identify the underlying complexity and inter-connectivity of molecular changes caused by CXCL4, we developed a new methodology (RegEnrich) to integrate the transcriptional and epigenetic layers and identify the important transcription factors modulated by CXCL4 (see Methods). Using RegEnrich, we first constructed weighted gene correlation networks, which allowed us to cluster genes into distinct modules (or sets of genes) based upon either their co-expression or co-methylation patterns (**Figure 3—figure supplement 2A-B**)(Langfelder & Horvath, 2008; Langfelder, Luo, Oldham, & Horvath, 2011). Modular analysis segregated the differentially expressed genes into 27 modules, each exhibiting a distinct co-expression pattern (**Figure 3B, C** and **Figure 3—figure supplement 2C**). Of these 4 were CXCL4-moDCs-specific modules (RM24-RM27) and contained genes belonging to ECM organization, ion channel transport, IFNα signaling and metabolic pathways, highlighting the impact of CXCL4 upon the DC phenotype (**Figure 3B** and **Figure 3—figure supplement 3B**). Similarly, modular analysis segregated all differentially methylated genes into 10 distinct co-methylation modules (**Figure 3B**). CXCL4-moDC-specific modules (DM5 and DM9) contained genes belonging to transcriptional and translational pathways, antigen presentation pathways, and the innate immune system (**Figure 3B** and **Figure 3—figure supplement 3C**; **Supplementary File 8**). However, we did not find much overlap between the CXCL4-moDC-specific co-expression and co-methylation modules (**Figure 3C**). Thus, DNA methylation only partially influences the transcriptional changes of CXCL4 signature genes.

To test whether transcription regulators are the central players (hubs) in our networks(Goode et al., 2016; Ramirez et al., 2017), we calculated module memberships as a measure to determine the importance of a gene in a given module(Langfelder & Horvath, 2008; Langfelder et al., 2011). Interestingly for both co-expression and co-methylation modules, we found that the transcription regulators typically exhibited higher module membership than the other genes (**Figure 3D, E** and **Figure 3—figure supplement 2D**). That transcription regulators are typically the hubs in our networks highlights their crucial regulatory function in modulating the expression dynamics of their downstream target genes. Thus, alterations in the expression and activity of a few key transcription regulators can potentially precipitate the large phenotypic differences observed between moDCs and CXCL4-moDCs. Using RegEnrich, we ranked the transcription regulators most prominently dysregulated between CXCL4-moDCs and conventional moDCs during differentiation (**Figure 4A**), and post stimulation (**Figure 4B**). Using these gene regulatory networks, we found that key transcription regulators such as *CIITA*, *TLE1*, *PTRF*, *MAPK13*, *CRABP2*, *IRF8*, regulate a large number of the CXCL4 signature genes, including pro-inflammatory and ECM pathway genes (**Figure 4A and B**). We confirmed that these key transcription regulators were significant for CXCL4 signature genes using two independent approaches: i) random forest-based gene regulatory networks (see Methods, and **Figure 4—figure supplement 1A-B**), and ii) literature-derived gene regulatory networks (data not shown). Together, the data-driven gene regulatory networks identified a direct mechanistic link between CXCL4, inflammation and ECM modeling in moDCs.

**Figure 4.**
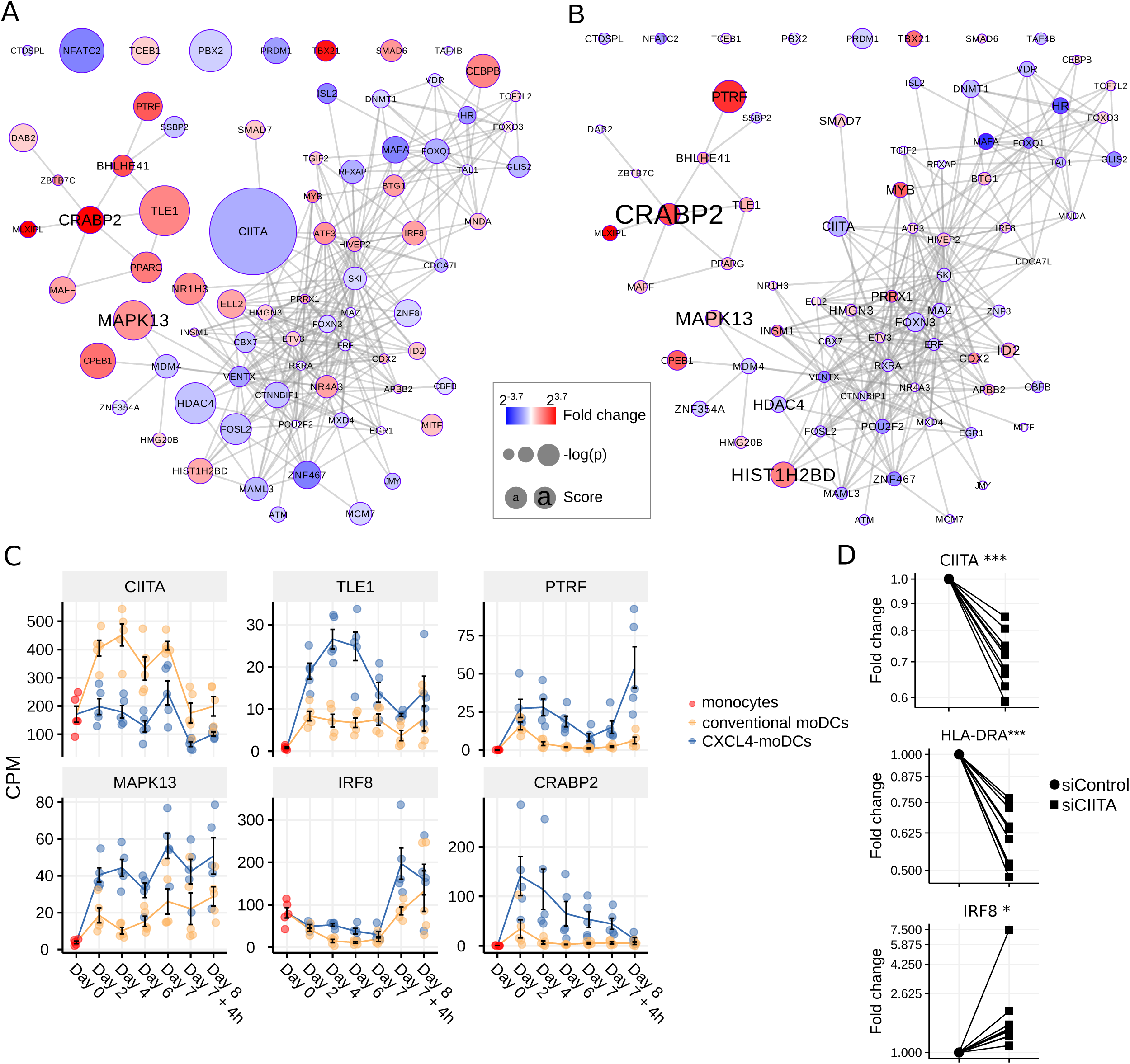
Transcription regulator enrichment highlights key TF candidates. Co-expression based key TF network during: **(A)** differentiation and **(B)** stimulation. Colors indicate fold change between CXCL4-moDCs and conventional moDCs. Red represents upregulation and blue represents downregulation. Circle size indicates −log_10_(p) for each comparison, where p is the p-value calculated during differential expression analysis; text size shows the RegEnrich score (see Methods). **(C)** Expression profile (mean±SEM) of key regulators. **(D)** Expression of *CIITA*, *HLADRA*, and *IRF8* on day 6 measured by qPCR in moDCs obtained from monocytes transfected with silencer negative control siRNA (siControl) or silencer CIITA siRNA (siCIITA). qPCR data were normalized using mean expression of *RPL32* and *RPL13A*. Fold change in y-axis (log2 scaled) is relative to the value obtained for siControl for each donor. Lines connect individual donors. \**P*<0.05, \*\*\**P*<0.001, paired two-sided Student’s *t*-test.

### CIITA is a key target of CXCL4 signaling

We found that the key transcriptional regulatory proteins exhibit different mRNA expression patterns over time. For example, *TLE1*, *PTRF* and *CRABP2* were expressed at low levels in monocytes but were upregulated during the differentiation of both conventional moDCs and CXCL4-moDCs (**Figure 4C**). However, these genes exhibited persistently higher expression in CXCL4-moDCs during both differentiation and following polyI:C stimulation (**Figure 4C**). Another example is the interferon regulatory factor 8 (IRF8), a transcription factor typically associated with pro-inflammatory gene expression in monocytic lineages, which is markedly upregulated in immature CXCL4-moDCs compared to conventional moDCs (**Figure 4C**). Class II MHC transactivator (CIITA), a transcription co-factor associated with regulation of MHC class II gene expression, was the most significantly down-regulated regulator in CXCL4-moDCs (**Figure 4C**).

To validate the inter-connectivity of important regulators inferred from the gene regulatory network, we performed siRNA-mediated knockdown to silence CIITA expression. At day 6 following introduction of siRNA, monocyte-derived cells remained viable, and displayed the anticipated phenotype: CIITA-silencing down-regulated expression of both CD74 and HLA-DR (**Figure 4D** and **Figure 4—figure supplement 1C-D**)(Landsverk, Gregers, & Bakke, 2012). While IRF8 has not been reported to be regulated by CIITA, our gene regulatory networks predicted direct or indirect regulatory interactions between CIITA and IRF8. Silencing of CIITA led to upregulation of IRF8, mimicking the effects of CXCL4 and validating the prediction of our gene regulatory networks (**Figure 4D** and **Figure 4—figure supplement 1C-D**). Hence using our gene regulatory networks, we have elucidated novel gene regulatory interactions in moDCs and found that CXCL4 alters the fate of moDCs by modulating the expression of the key transcription regulator CIITA.

### CXCL4 induces fibrotic pathways in moDCs mediated via epigenetic imprinting and CIITA

Our data-driven methodology allowed us to identify several novel regulators and pathways that are differentially regulated due to CXCL4 during moDC differentiation (**Figure 2F**, **4A-C** and **Supplementary File 7**). As a result, we observed that even unstimulated CXCL4-moDCs exhibit a pro-fibrotic phenotype, as characterized by the increased gene and protein expression of several crucial ECM-related molecules including FN1, SPP1, IL1RN and TGFB1 (for transcriptional changes see **Figure 5A**, **Figure 5—figure supplement 1A-B**; for protein validations see **Figure 5B-D**). Importantly, silencing CIITA mimicked the effects of CXCL4 leading to upregulation of FN1 along with other molecules involved in ECM remodeling validating the relevance of this network in the pro-fibrotic cascade (**Figure 5E-G**, **Figure 5—figure supplement 1C-D**). Since in CXCL4-moDCs we found that majority of the differential genes were hypermethylated (**Figure 2A and B**) and that ECM genes were up-regulated (including FN1 and TGFB1; see **Figure 5A and B**), we next examined whether modulating DNA methylation affects FN1 and TGFB1 expression. In line with our hypothesis, inhibition of DNMTs using 100nM 5-Aza-2′-deoxycytidine restored the expression of FN1 and TGFB1 which were upregulated by CXCL4 in moDCs (**Figure 5H**), suggesting that CXCL4 associated epigenetic imprinting also plays a role in promoting expression of pro-fibrotic genes. Although our data provides unprecedented data for the direct implication of CXCL4 in tissue fibrogenesis via CXCL4-moDCs, we next examined the possible implication of these CXCL4-moDCs on fibroblast behavior. By culturing fibroblasts with the supernatant of CXCL4-moDCs stimulated with polyI:C, we demonstrate that these fibroblasts expressed markedly higher levels of inflammatory mediators associated with fibrosis and above all, myofibroblast transition, considered indispensable for fibrosis, compared to conventional moDCs stimulated with the same TLR3 ligand (**Figure 5I** and **Figure 5—figure supplement 2**). Together, this data unequivocally demonstrates that CXCL4 alters the fate of moDCs differentiation into cells that drive fibrogenesis both directly and, via myofibroblast activation, indirectly.

**Figure 5.**
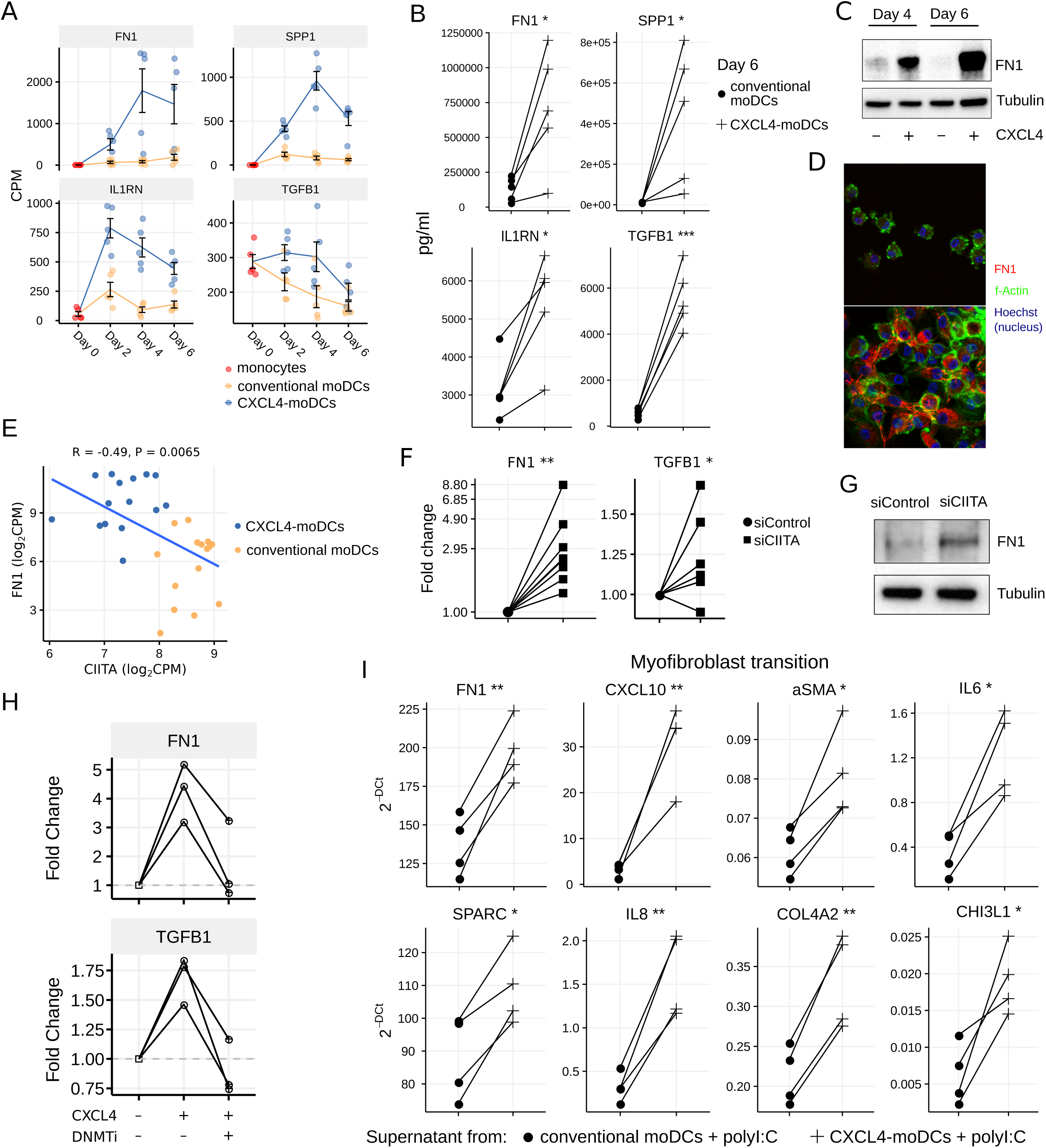
CXCL4 induces production of ECM components in moDCs and fibroblasts. **(A)** Expression of genes implicated in ECM remodeling (mean±SEM). **(B)** Validation (luminex) of ECM protein production in CXCL4-moDCs and conventional moDCs on day 6. **(C)** Fibronectin (FN1) expression (tubulin as loading control) determined using Western blot on days 4 and 6 (representative of 5 independent experiments). **(D)** Fibronectin (red) synthesis determined using confocal imaging on day 6 (green: f-actin; and blue: nucleus staining using Hoechst). **(E)** Pearson correlation between gene expression of *CIITA* and *FN1* during differentiation (i.e. on day 2, 4 and 6). **(F)** FN1 and TGFB1 expression measured by qPCR and **(G)** FN1 expression measured by western blot on day 6 moDCs obtained from monocytes transfected with siControl and siCIITA (see Figure 4D). **(H)** FN1 and TGFB1 expression measured by qPCR on day 3 in conventional moDCs, CXCL4-moDCs and CXCL4-moDCs exposed to DNMT inhibitor (100nM 5-Aza-2′-deoxycytidine). **(I)** Expression of ECM genes measured using qPCR in healthy dermal fibroblasts (one representative donor; for others see **Figure 5—figure supplement 2**) co-cultured with supernatants from CXCL4-moDCs and moDCs that were stimulated for 24 hours with polyI:C. qPCR data were normalized using mean expression of *RPL32* and *RPL13A*. In panels **B**, **F**, **H** and **I**, lines connect individual donors. \**P*<0.05; \*\**P*<0.01, \*\*\**P*<0.001, paired two-sided Student’s *t*-test.

## Discussion

Although the role of inflammation in fibrosis is increasingly recognized, the underlying molecular links between these processes remain elusive and their identification is paramount for the development of medicines to not only halt progression but prevent fibrosis. Using whole genome transcriptional and epigenetic profiling, we find that CXCL4 drives the development of a pro-inflammatory and pro-fibrotic phenotype in moDCs, characterized by the excessive production of ECM components and capacity to promote myofibroblast differentiation. As these are two key mechanisms contributing to tissue fibrogenesis, our study introduces the novel concept that CXCL4-induced inflammatory moDCs constitute the driving force behind both the initiation and progression of fibrosis in diseases where CXCL4 levels are increased such as SSc.

TGF-β is considered a key regulator during fibrosis in physiological and pathological conditions(Massague, 2012). For instance, TGF-β drives mesenchymal responses during wound healing, where its transiently increased expression promotes myofibroblast transition. However, the initial stage of wound healing is the formation of a platelet plug, followed by monocyte recruitment and monocyte differentiation into M1 macrophages. After this primarily inflammatory phase, a switch to resolution, accompanied by tissue repair and fibrosis, occurs(Xue & Jackson, 2015). Platelets, crucial players in the pathogenesis of several diseases including SSc, contain large amounts of CXCL4(Levine & Wohl, 1976; van Bon, Affandi, et al., 2014; van Bon et al., 2016). Activation of platelets early on in the wound healing process is likely to precede the synthesis and secretion of TGF-β. Notably, CXCL4 was found to play an important role in lung inflammation and tissue damage(Hwaiz, Rahman, Zhang, & Thorlacius, 2015), and has been identified as a biomarker for early rheumatoid arthritis where it was co-localized with inflammatory cells and platelets in synovial tissue(Yeo et al., 2016). In contrast to other inflammatory mediators that appear at later stages of disease, CXCL4 levels are also increased in patients at risk for SSc, a disease in which clinical inflammation precedes fibrosis by years(van Bon, Affandi, et al., 2014). Together, these observations indicate an early role for CXCL4 in inflammatory and subsequent fibrotic processes, placing CXCL4 upstream of TGF-β. This possibility is further substantiated by our finding that CXCL4 clearly induces TGF-β RNA and protein expression (**Figure 5A and B**).

Multiple studies provide compelling evidence for the presence of inappropriately activated and/or trained innate immunity in patients with inflammatory diseases. Recently, several crucial studies have highlighted the molecular basis, relevance and pathological consequences of innate immunity trained by various exogenous ligands and endogenous ligands the latter contributing to atherosclerosis and gout(Bekkering et al., 2014; Cheng et al., 2014; Crisan et al., 2016; Mourits, Wijkmans, Joosten, & Netea, 2018). Following a seminal study which observed enhanced collagen synthesis in SSc patient skin fibroblasts compared to those of healthy control(LeRoy, 1974), this phenomenon was observed in SSc patient DCs, which had potentiated responses to various TLR agonists(van Bon, Affandi, et al., 2014; van Bon, Cossu, et al., 2014). Our study now reveals that differentiating monocytes undergo massive transcriptomic and epigenetic reprogramming upon CXCL4 exposure, and we propose that CXCL4 is a clinically relevant and important endogenous ligand bridging inflammation with fibrosis via trained immunity and provides a rationale for therapeutic targeting of CXCL4 in SSc and other fibrotic diseases.

## Materials and Methods

### Differentiation and stimulation of CXCL4 moDCs

Blood from healthy donors (HDs) was collected in accordance with institutional ethical approval. Peripheral blood mononuclear cell (PBMC) and monocyte isolation, as well as differentiation of monocyte-derived dendritic cells (moDCs) were performed as described previously(Silva-Cardoso et al., 2017). Briefly, PBMCs were isolated from heparinized venous blood using Ficoll Paque^TM^ Plus (GE Healthcare) density gradient. Monocytes were purified with anti-CD14 magnetic beads-based positive isolation using autoMACS Pro Separator-assisted cell sorting (Miltenyi Biotec), according to the manufacturer’s protocol. Monocyte purity was above 95% for all of the samples (data not shown). For the differentiation of moDCs, monocytes were cultured at a density of 1×10^6^ cells/ml in culture medium comprised of RPMI 1640 with GlutaMAX (Life Technologies) supplemented with 10% (v/v) heat-inactivated fetal bovine serum (FBS; Biowest) and 1% (v/v) antibiotics (penicillin and streptomycin; Life Technologies). In order to generate moDCs, GM-CSF (800 U/ml; R&D) and IL-4 (500 U/ml; R&D) were added. For the experiments where we investigated the effects of CXCL4, we added 10 µg/ml of recombinant human CXCL4 (PeproTech) on day 0 and day 3. Medium and cytokines were refreshed on day 3. Differentiated moDCs were obtained after 6 days from monocytes cultured at 37°C in the presence of 5% CO_2._ After differentiation, cells were washed, plated at a density of 0.5×10^6^ cells/ml and left overnight (O/N) in new culture medium. Cells were stimulated with 25 µg/ml of polyinosinic-polycytidylic acid (polyI:C; InvivoGen) for 4 hours or 24 hours, or kept unstimulated, as shown in **Figure 1A**.

### 5-aza-2’-deoxycytidine treatment

Monocytes were seeded at a density of 1×10^6^ cells/ml and cultured in medium supplemented with IL-4, GM-CSF with (for conventional moDCs) or without CXCL4 (for CXCL4-moDCs), as described above. Cells differentiating into CXCL4-moDCs were either left untreated or were treated with the DNA methyltransferase inhibitor 5-Aza-2′-deoxycytidine (Sigma Aldrich) at the concentration of 100nM. On day 3, cells were harvested and processed for RNA analysis.

### DNA and RNA extraction for DNA methylation and RNA sequencing analysis

For DNA methylation and RNA sequencing analysis cells were collected from 5 HDs: on the first day of culture (monocytes, day 0); during differentiation on day 2, day 4 and day 6. After O/N resting, unstimulated cells (day 7), cells stimulated with polyI:C for 4 hours (day 7 + 4h) and 24 hours (day 8) were also lysed in RLTplus buffer (Qiagen) containing 1% (v/v) beta-mercaptoethanol (Sigma). In total, we obtained 65 paired samples for RNA sequencing and DNA methylation profiling. DNA and RNA were extracted using the Allprep Universal Kit (Qiagen) following the manufacturer’s instructions. For the experimental validation using transfected moDCs and fibroblasts, due to the limiting number of cells, the total RNA was isolated using an RNeasy Micro Kit (Qiagen) according to the manufacturer’s instructions. The concentration of DNA and RNA was assessed using the Qubit RNA HS Assay Kit and Qubit dsDNA HS Assay Kit (Life Technologies), respectively, and measured in the Qubit 2.0 fluorimeter (Invitrogen).

### RNA sequencing

RNA Sequencing (RNA-seq) was performed at the Genomic Facility from the University Medical Center of Utrecht. RNA integrity was first evaluated using a Bioanalyzer (Agilent). RNA-seq library was prepared using 100ng total RNA using the TruSeq kit (Illumina). Oligo(dT) magnetic beads were used to enrich for messenger RNAs which were then fragmented (about 200 bp). Random hexamer-primers were used to reverse transcribe mRNA into double stranded cDNA, which was then end-repaired followed by addition of 3’-end single nucleotide adenine. Sequencing adaptors were ligated to the resulting cDNA that was subsequently amplified using PCR. Agilent 2100 Bioanaylzer and the ABI StepOnePlus Real-Time PCR System were used to assess the quality and quantity of RNA-seq libraries. The library products were sequenced on an Illumina NextSeq 500 sequencer using 75bp single-end reads, generating on average 26.2 million clean reads per sample.

### Transcriptional data analysis

For each of the 65 transcriptional profiles, reads were aligned using STAR aligner using the default parameters to the 65,217 annotated genes obtained from the GrCh38 (v79) built from the human genome (http://www.ensembl.org). On average 22.5 million uniquely mapped reads were obtained per sample. The read counts per gene were quantified by the Python package HTSeq(Anders, Pyl, & Huber, 2015) using annotations from the GrCh38 (v79) built from the human genome (http://www.ensembl.org). Differentially expressed genes (DEGs) were identified by using the DESeq2 (1.8.2) Bioconductor/R package(Love, Huber, & Anders, 2014) using likelihood ratio test (LRT), and genes with FDR adjusted p-value < 0.05 were considered differentially expressed. Raw count data were transformed to count per million (CPM) for gene expression visualization. Variance stabilizing transformation (VST) was applied to obtain the VSD data for further analysis(Love et al., 2014).

### DNA methylation profiling

DNA methylation profiling was performed at the GenomeScan (GenomeScan B.V., Leiden, The Netherlands). Genomic DNA was bisulfite-converted using the EZ DNA Methylation Gold Kit (Zymo Research) and used for microarray-based DNA methylation analysis on the HumanMethylation850 BeadChip (Illumina, Inc.), according to the manufacturer’s instructions. Beadchip images were scanned on the iScan system and the data quality was assessed using the minfi (version 1.20.2) package(Aryee et al., 2014) using default analysis settings.

### DNA methylation data analysis

Illumina Infinium HumanMethylation850 BeadChip fluorescent data (>850,000 CpG sites) were imported and transformed to methylated (M) and unmethylated (U) signal by minfi package(Aryee et al., 2014). CpG probes were quality-checked and filtered using the following criteria: (i) probes that failed in at least 5% samples were removed, (ii) probes with bead count < 3 in at least 5% of samples were removed, (iii) probes targeting SNP sites were removed, and (iv) probes that aligned to multiple locations were removed as described(Nordlund et al., 2013). We further removed the probes for the sex chromosomes. One sample (102920-001-17, moDC differentiation sample from donor 4 on day 2) did not pass the quality check and was removed from the subsequent analysis. Approximately 558,000 CpG sites located in six regions (TSS1500, TSS200, 5’UTR, 1stExon, Exon boundaries and 3’UTR) remained after the quality checks. The intra-array data normalization for the bias introduced by two types of Infinium probes was performed by Beta-mixture quantile normalization (BMIQ) method in ChAMP (version 2.6.0) package(Morris et al., 2014). The DNA methylation level of each CpG was depicted by the ratio of methylated (M) signal relative to the sum of both methylated and unmethylated (U) signal:

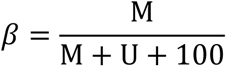

We studied the alterations of DNA methylation considering: i) individual CpG sites, ii) region of the CpG site (including 1500 base pairs before TSS or TSS1500, TSS200, 5’UTR, 1st Exon, Exon boundaries and 3’UTR), and iii) proximal genes. To find the differentially methylated CpGs (DMPs) of moDC or of CXCL4 moDC associated with time, a linear regression model with two variables (donor and time) was fitted at each probe. We analyzed DMPs separately for differentiation and stimulation experiments. CpG sites with time-associated FDR corrected p-value <0.05 were considered DMPs. Similarly, DMPs between moDCs and CXCL moDCs were identified using a linear regression model with three variables (donor, time and condition). To obtain region-specific β-values, we calculated the average β-values using all probes that mapped to the same region (including TSS1500, TSS200, 5’UTR, 1stExon, ExonBnd and 3’UTR(Jiao, Widschwendter, & Teschendorff, 2014)) for a given gene. We then applied the same regression models to find differentially methylated regions (DMRs). If any of the regions around the gene were significantly altered, we considered that the gene was differentially methylated (DMGs).

### Multidimensional scaling (MDS) analysis

Transcriptional data (VSD) and DNA methylation data (β) were utilized to visualize the differences of cells during differentiation and polyI:C stimulation. The Euclidean distances between samples were calculated based on VSD or β. Multidimensional scaling was performed using these distances in R (cmdscale function from stats package) to project (visualized using ggplot2 package) the high dimensional transcriptional or DNA methylation data onto two dimensions. MDS plots were generated using the DEGs or DMGs.

### Comparison of gene expression and DNA methylation

To compare the relationship between expression and methylation data, we analyzed the two-layered data from genes that were both differentially expressed and methylated. We calculated Spearman correlation coefficients (for **Figure 3A**, and **Figure 3—figure supplement 1B-D**) between the expression (VSD) and methylation (β values) data for the genes using the cor function in R. To ensure paired analysis, we removed the corresponding expression profiles for the sample which failed the DNA methylation quality checks. Thus, we performed all correlation-based analysis using 64 pairs of samples. To study the global relationship between gene expression and DNA methylation, contour plots were constructed for paired expression (VSD) and methylation (β values) data using geom_density2d function in R (**Figure 3—figure supplement 1F and G**). For **Figure 3—figure supplement 1F** we used the paired data from all the genes, while for **Figure 3—figure supplement 1G** we used the paired data from all the genes that were both differentially expressed and differentially methylated. We further analyzed the relationship between the paired expression (VSD) and methylation (β values) data using linear regression models and by fitting smoothing curves using generalized additive model (GAM).

### Pathway enrichment analysis

Pathway enrichment analysis, for DEGs, DMGs or module genes, was performed using hypergeometric test in ReactomePA package(Yu & He, 2016). The compareCluster function in the ReactomePA package (with parameters fun=“enrichPathway”, pAdjustMethod = “fdr”, and pvalueCutoff = 0.05) was used to compare and plot the pathways enriched in different sets of genes.

### CIITA-silencing in monocytes

Freshly purified monocytes were cultured in medium without antibiotics at a density of 2×10^6^ cells/ml. Transfection mix was prepared with 40nM of Silencer^®^ pre-designed siRNA against human CIITA (targeting exon 3 and 4; siCIITA) or the Silencer^TM^ Negative Control No.1 (siControl) (Life technologies), Lipofectamine 2000 and Plus Reagent (both from Invitrogen), diluted in Opti-MEM^®^ I Reduced-Serum Medium (Life Technologies). After 5 hours, transfected cells were washed with culture medium and were differentiated into moDCs as described above.

### Fibroblast cultures

Dermal fibroblasts (DF) were isolated from healthy skin biopsies. Skin biopsies were obtained from unused material after cosmetic surgery from anonymous donors who had given prior informed consent to use the biopsies for research. The use of this material is exempted from ethical review processes. DF were isolated using the Whole Skin Dissociation Kit (MiltenyiBiotec) following the manufacturer’s instructions, cultured in DMEM medium (Life Technologies) supplemented with 10% (v/v) FBS, and 1% (v/v) antibiotics (used for experiments between passages 4 and 5). Prior to the treatment, DF were cultured O/N with DMEM medium containing 1% FBS. Supernatants collected from moDCs and CXCL4 moDCs stimulated with polyI:C for 24 hours were added to the DF for 24 hours. Medium and polyI:C were also added to the DF as controls (data not shown).

### Real-time quantitative PCR

Purified RNA was retro-transcribed with iScript Reverse Transcriptase Kit (Bio-Rad). Gene expression was measured by Real-Time quantitative-PCR (RT-qPCR) on the QuantStudio 12k flex system using SybrSelect Mastermix (Life Technologies). To calculate the ratio between the expression of a gene of interest and housekeeping genes (mean between *RPL32* and *RPL13A* for moDCs cultures; RPL13A for the fibroblasts cultures), we used either the 2^-DCt^ or the 2^-DDCt^ method. Primer sequences are listed in the online **Supplementary File 2**.

### Cytokine production measurement

To validate secreted targets at the protein level, we collected cell-free supernatants after moDC and CXCL4 moDCs differentiation (day 6) and after 24 hour stimulation with polyI:C (day 8) from the same 5 HDs that were used for RNA sequencing and DNA methylation profiling. Cytokine measurements were assessed using Luminex assay as previously described(Silva-Cardoso et al., 2017) at the MultiPlex Core Facility of the Laboratory of Translational Immunology (University Medical Center Utrecht). Data were acquired using Bio-Rad FlexMap3D system and the Xponent 4.2 software, and analyzed using Bio-Plex Manager (version 6.1).

### Flow cytometry

Cells were first incubated with Fixable Viability Dye eFluor780 (eBioscience) in PBS to exclude dead cells and were further treated with 10% (v/v) mouse serum (Fitzgerald). Next, cells were stained with the following anti-human fluorochrome-conjugated mAbs: CD14 (clone M5E2), CD86 (clone IT2.2) and CLEC10A (H037G3) obtained from BioLegend, CD1a (clone HI149) and LAMP1/ CD107a (clone H4A3) obtained from BD. Cells were acquired on the LSR Fortessa (BD) and data was analyzed using the FlowJo software (version 7.6.5; Tree Star. Inc.).

### Western blot

Cells were washed with PBS and lysed in Laemmli buffer. Protein concentration was quantified using the Pierce BCA Protein Assay Kit (Thermo Scientific) according to the manufacture’s protocol. Equal amounts of protein from different lysates were separated by electrophoresis on a 4-12% Bis-Tris SDS NuPAGE gels (Invitrogen) and transferred to a PVDF membrane (Millipore). After blocking the membranes with Tris-buffered saline (pH 8) containing 0.05% Tween-20 and 4% milk (Bio-Rad) for 1 hour at room temperature (RT), the membranes were probed with the antibodies recognizing FN1 (Abcam) and tubulin (Sigma-Aldrich) O/N at 4°C. Afterwards, membranes were washed and incubated for 1 hour at RT with the secondary anti-rabbit or anti-mouse antibodies, both HRP-conjugated (Dako). Protein detection was assessed using a ChemiDoc MP System (Bio-Rad). Protein visualization and densitometry analysis of band intensity were performed using the Image Lab software (version 5.1, Bio-Rad). We calculated the ratio between the expression of FN1 and tubulin to determine the relative expression of FN1 in different conditions.

### Confocal microscopy

As an alternative way to validate the expression and production of FN1, we performed microscopy analyses as described before(Silva-Cardoso et al., 2017), with minor modifications. For the differentiation of moDCs, we used Nunc^®^ Lab-Tek^®^ II chamber slides (Thermo Scientific) pre-coated with 0.01% (v/v) poly-L-lysine (Sigma-Aldrich) in sterile water. After differentiation, cells were incubated with fixation/permeabilization solution (eBioscience) supplemented with 5% (v/v) normal goat serum (Cell Signaling) for 30 minutes at RT, followed by two washes with permeabilization buffer (eBioscience). Cells were incubated for 1 hour with primary antibody recognizing FN1 (Abcam). After washing twice, cells were incubated with secondary antibody Alexa 594 anti-rabbit (Life Technologies) and phalloidin-labeled FITC (ENZO) for 1 hour. Cells were washed and incubated with Hoechst 33342 (1 µM; Invitrogen) for 15 minutes. Next, cells were washed with permeabilization buffer twice, and at last washed with 1% (w/v) BSA and 0.1% (v/v) sodium azide (NaN_3_; Sigma-Aldrich) in PBS. Mowiol (Sigma-Aldrich) was used to mount the dry slides and coverslips. Image acquisition was performed on a LSM710 (Zeiss) confocal microscope using the Zen2009 (Zeiss Enhanced Navigation) acquisition software. Confocal images were obtained with the objective 63x 1.40 oil and analyzed using the ImageJ software.

## RegEnrich pipeline

We developed a data driven pipeline (RegEnrich) to integrate CXCL4 specific transcriptional and DNA methylation signatures and to predict the key TFs driving the differential transcriptional profile of CXCL4 moDCs compared to moDCs (**Figure 3B-E**, and **4A-B**). RegEnrich pipeline involves three steps: 1) construction of data-driven networks; 2) deducing genes of interest; and 3) enrichment of transcriptional factors or regulators (henceforth called “*TF*”). The aim of RegEnrich pipeline is to rank TFs based on their differential expression and the enrichment of their own downstream targets in a given gene set. R functions and codes used for performing the data analysis are available at: https://bitbucket.org/systemsimmunology/cxcl4_modcs

### Co-expression/co-methylation network construction

For co-expression network, VSD data of all DEGs were used to construct a co-expression network by R package WGCNA (version 1.51)(Langfelder & Horvath, 2008) as described in https://labs.genetics.ucla.edu/horvath/CoexpressionNetwork. Briefly, we used unsigned correlations and a soft thresholding power of 6 to construct networks with scale free topology. We calculated the adjacency matrix which was further used to calculate Topological overlap matrix (TOM) to identify modules of co-expressed genes. Modules were identified using cutreeDynamic function with the minimum module size of 30. Modules were further merged if the Pearson correlation of their eigengene was <0.25. Using this methodology, we obtained 27 co-expression modules (**Figure 3—figure supplement 2A**). Nodes (genes) and edges (connections of genes) in each module were exported by exportNetworkToCytoscape function (threshold >=0.02).

To build co-methylation network, we first assigned a unique β value to a given gene of a sample by setting priority to four regions: TSS200>TSS1500>5’UTR>1stExon as described in Jiao *et al*.(Jiao et al., 2014). If for a gene TSS200 region is differentially methylated (DM), we considered the β value of TSS200 as the methylation level of this gene. Similarly, for a gene without DM TSS200 but with DM TSS1500, β value of TSS1500 region was used, and so on. Then these regions, representing corresponding genes, were used to build co-methylation networks using the methodology described for the co-expression network. To achieve topological scale-free networks, standard parameters were set to a soft thresholding power of 12, “unsigned” network, minimum module size of 30, merged module threshold < 0.25, and an exporting network threshold of 0.02. In total, 10 modules were reserved in the end (**Figure 3— figure supplement 2B**).

### Integration of co-expression and co-methylation network

Spearman correlation coefficients were calculated using the co-expression and co-methylation module eigengenes to integrate the two networks (**Figure 3—figure supplement 2C**). We calculated the number of genes shared between co-expression and co-methylation modules and two-tailed fisher exact test was used to evaluate the significance of each overlap (**Figure 3C**). Pearson correlation coefficients were used to relate gene modules to sample traits i.e., CXCL4^+/-^, time and polyI:C^+/-^ (**Figure 3C**).

### Gene regulatory network (GRN) construction based on random forest (RF) algorithm

To obtain potential transcription (co-)factor/regulators (TF) for each gene in a data-driven manner, we constructed a TF-target GRN using random forest machine learning algorithm (modified from(Huynh-Thu, Irrthum, Wehenkel, & Geurts, 2010; Walley et al., 2016) and is part of RegEnrich package developed by us; see below). This TF-target GRN is a directed network of two types of components: a) TFs and b) their potential targets. Here targets might not necessarily be direct downstream targets that the TFs might bind to, rather the genes that are inferred to be directly/indirectly regulated by the TFs based on the transcriptional data. The construction of TF-target GRN consisted of four steps. First, the VST normalized data from 17,709 DEGs (same genes in co-expression network), including 1,172 differentially expressed TFs (**Supplementary File 9**), were selected for the analyses. Second, we removed all the target genes that are expressed in less than 10 samples. Third, for every target gene, a random forest model was built to predict its expression based on TF expression (the parameters are: K = “sqrt”, nb.trees = 1000, importance.measure = “IncNodePurity”). As a last step, models with low performance (MSE < 0.5) were removed to achieve a robust TF-target GRN.

### TF enrichment analysis

In this study, TF enrichment analyses were performed on two data-driven networks, a co-expression network and a GRN network. For the co-expression network, we ranked the edges between TFs and their potential targets based on the edge weight. Top 5% edges were then selected and were considered for further analyses. This resulted in 1,037,689 TF-target connections. Similarly for the GRN network, we used top 5% of edges (688,559 TF-target connections). One-tailed hypergeometric test was used to calculate the enrichment p-values (*P_E_*) for each TF in a given set of genes (here genes differential between CXCL4 moDCs and moDCs). Those TFs that exhibited significant differential expression (*P_D_* < 0.05) and had significant enrichment (*P_E_* < 0.05) were considered as key TFs. In other words, TFs that were differentially expressed along with their own targets were considered to be enriched in a given gene set. The overall scores of TFs were calculated by:

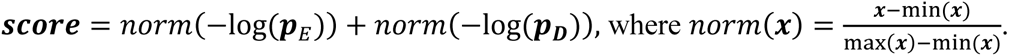

Cytoscape 3.4 (www.cytoscape.org) was used to visualize the networks. In TF-TF networks, we only plot the edges connecting the enriched key TFs in both co-expression and GRN network. For better visualization, only TFs with |log_2_(fold change)| > 0.6 were shown (**Figure 4A, B**, **Figure 4—figure supplement 1A-B**).

## Data availability

RNAseq count matrices and BAM files have been deposited in the National Center for Biotechnology Information’s Gene Expression Omnibus under accession number GSE115488. Raw and processed DNA methylation data has been deposited in the National Center for Biotechnology Information’s Gene Expression Omnibus under accession number GSE115201.

## Acknowledgments

We thank Dr. Kris Reedquist and Prof. Linde Meyaard for their critical comments, Dr. Marzia Rossato for discussions about RNAseq and other members of our lab for the fruitful discussions.

## Author contributions

Conceptualization (TRDJR, MB, WT, AP), Methodology (SCSC, BG, MC, WT, CA, APL, CPJB), Formal Analysis (WT, AD, AP), Resources (TRDJR), Writing original draft (SCSC, WT, AP, TRDJR), Writing reviewing and editing (SCSC, WT, CA, APL, CPJB, AD, JL, WM, EH, RJB, MB, AP, TRDJR), Visualization (WT, SCSC, AP), Supervision (TRDJR, MB, AP), Project Administration (TRDJR).

## Supplementary Figures

**Figure 1—figure supplement 1.**
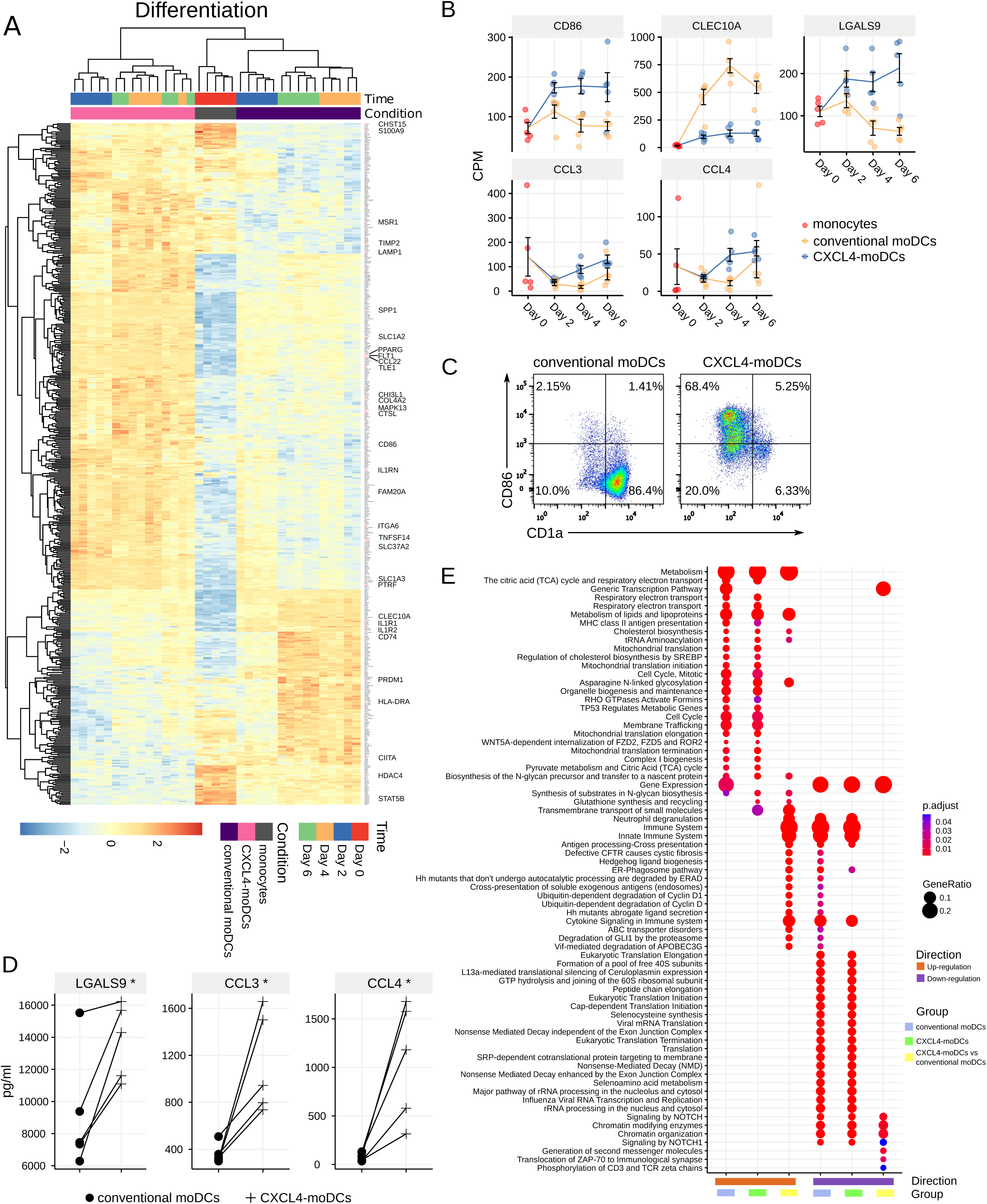
Differentially expressed genes during differentiation. **(A)** Heatmap showing normalized expression (VSD) of top 500 DEGs between CXCL4 moDCs and moDCs during their differentiation (day 0 to day 6). Heatmap color schemes are based on z-scores. Hierarchical clustering dendrograms were calculated using Euclidean distance. **(B)** Gene expression profiles for example genes from top 500 DEGs that were further validated at the protein level. Data are shown as mean of count per million (CPM) ± SEM. **(C)** Flow cytometry dot plot showing expression of CD86 and CD1a. Representative data from 5 HDs are shown. **(D)** Cytokine production of selected example proteins measured by Luminex using cell-free supernatants collected on day 6. Each symbol represents an individual donor; lines connect the same donor. **(E)** Pathway enrichment analysis for the genes which were significantly up- and down-regulated during differentiation, as shown in Figure 1B. Up to 30 most significant pathways (FDR adjusted p < 0.05) were shown for each set of genes. The size of the circle depicts the gene ratio of DEGs in the pathway to the total number of DEGs in each set of genes. Circle color represents FDR adjusted p values. The full lists of pathways are available in **Supplementary File 5**. Paired two-sided Student’s *t*-test; * *P*<0.05.

**Figure 1—figure supplement 2.**
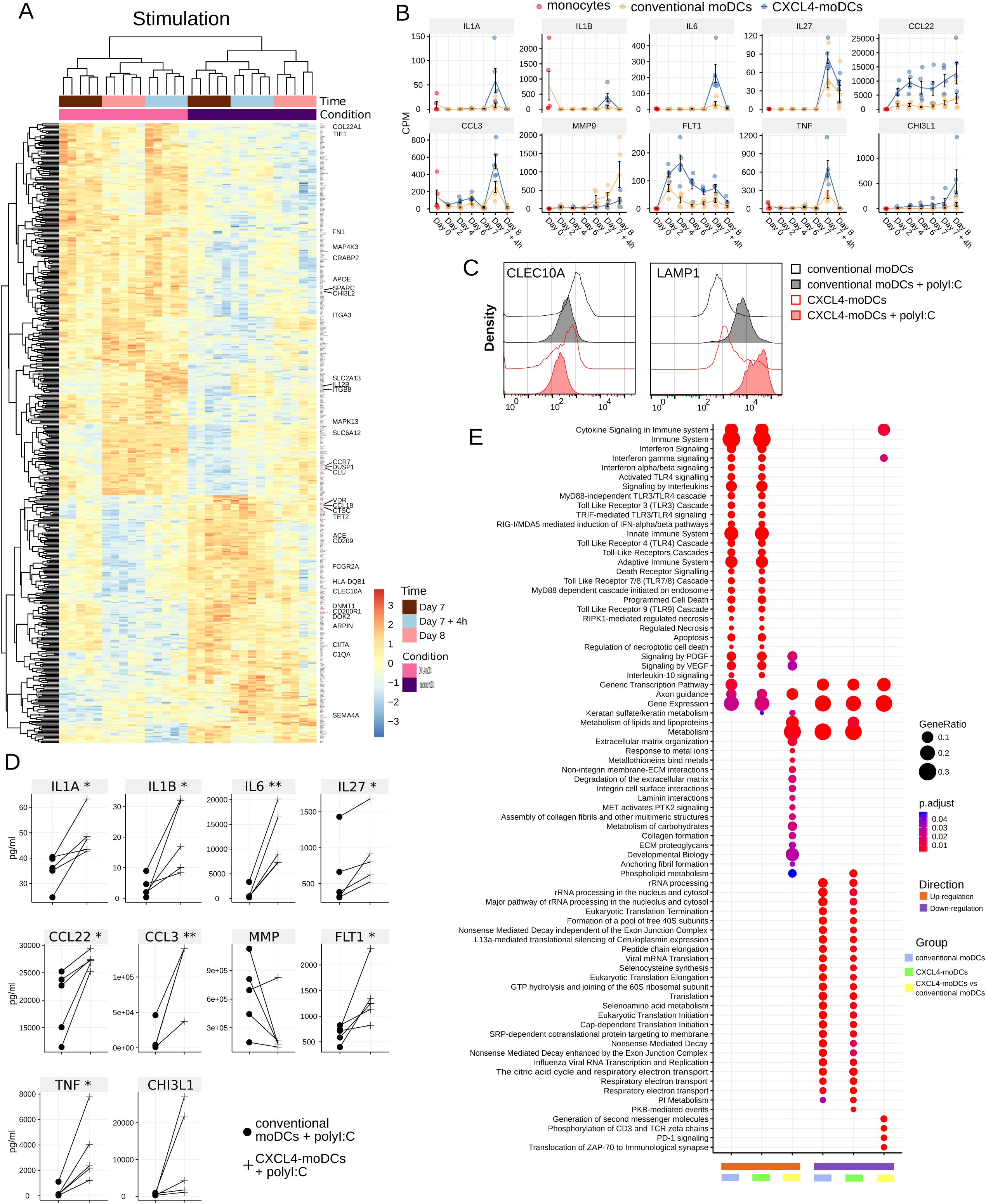
Differentially expressed genes upon polyI:C stimulation. **(A)** Heatmap same as **Figure 1—figure supplement 1** for top 500 DEGs between CXCL4 moDCs and moDCs upon polyI:C stimulation (day 7 to day 8). **(B)** Gene expression profiles for example genes from top 500 DEGs that were further validated on protein level. Data are shown as mean of count per million (CPM) ± SEM. **(C)** Flow cytometry analysis showing the relative expression of CLEC10A and LAMP1 between CXCL4 moDCs and moDCs, unstimulated and stimulated with polyI:C for 24 hours (on day 8). Representative data from 5 HDs are shown. **(D)** Cytokine production was measured by Luminex on cell-free supernatants after 24 hours stimulation with polyI:C (day 8). Each symbol represents an individual donor; lines connect the same donor. **(E)** Pathway enrichment analysis for the genes that were significantly up- and down-regulated upon polyI:C stimulation, shown in Figure 1C. Up to 30 most significant pathways (FDR adjusted p < 0.05) were shown for each set of genes. The size of the circle depicts the gene ratio of DEGs in the pathway to the total number of DEGs in each set of genes. Circle color represents FDR adjusted p values. The full lists of pathways are available in **Supplementary File 6**. Paired two-sided Student’s *t*-test; * *P*<0.05, ** *P* < 0.01.

**Figure 2—figure supplement 1.**
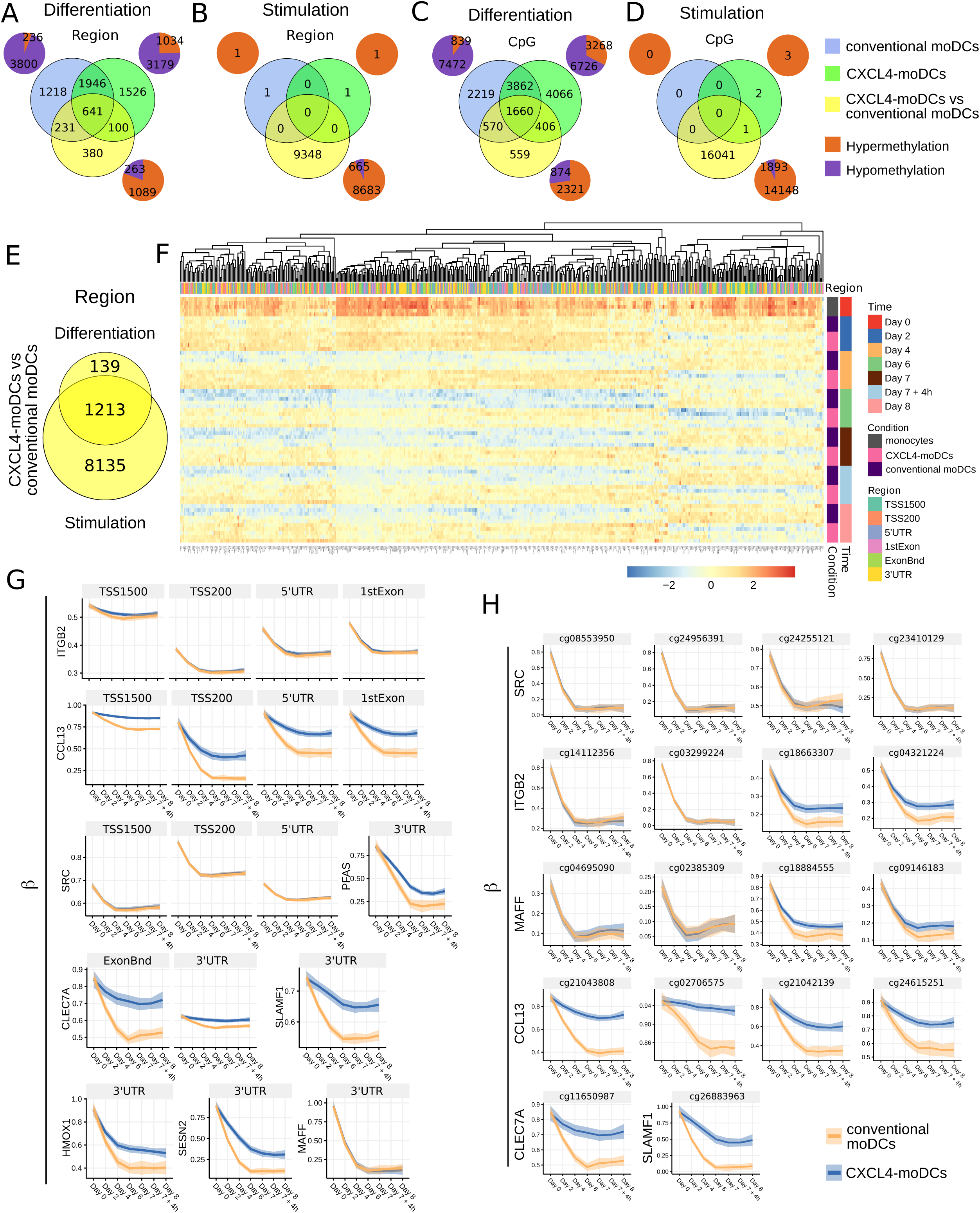
Dynamics of DNA methylation on region and CpG levels. **(A)** Venn diagram shows the overlaps of differentially methylated regions (DMRs) and CpG sites **(C)** during differentiation of: monocytes into moDCs (blue); monocytes into CXCL4 moDCs (green), and DMRs between moDCs and CXCL4 moDCs (yellow). (**B**) Venn diagram showing DMRs and CpG sites **(D)** in moDC (blue), CXCL4 moDCs (green) and DMR/CpG between moDCs and CXCL4 moDCs (yellow) after polyI:C stimulation. In **(A-D)** pie charts represent the number of hyper-methylated regions (orange) and hypo-methylated regions (magenta). **(E)** Venn diagram showing the overlap of DMRs during differentiation (yellow circle in panel **A**) and upon polyI:C stimulation (yellow circle in panel **B**). **(F)** Heatmap reporting top 500 regions in overlapping DMRs in panel **D**. **(G)** Temporal methylation patterns of selected regions and CpG sites **(H)** found to be differential in different comparisons. Lines represent mean of β values and shades represent 95% confidence interval.

**Figure 3—figure supplement 1.**
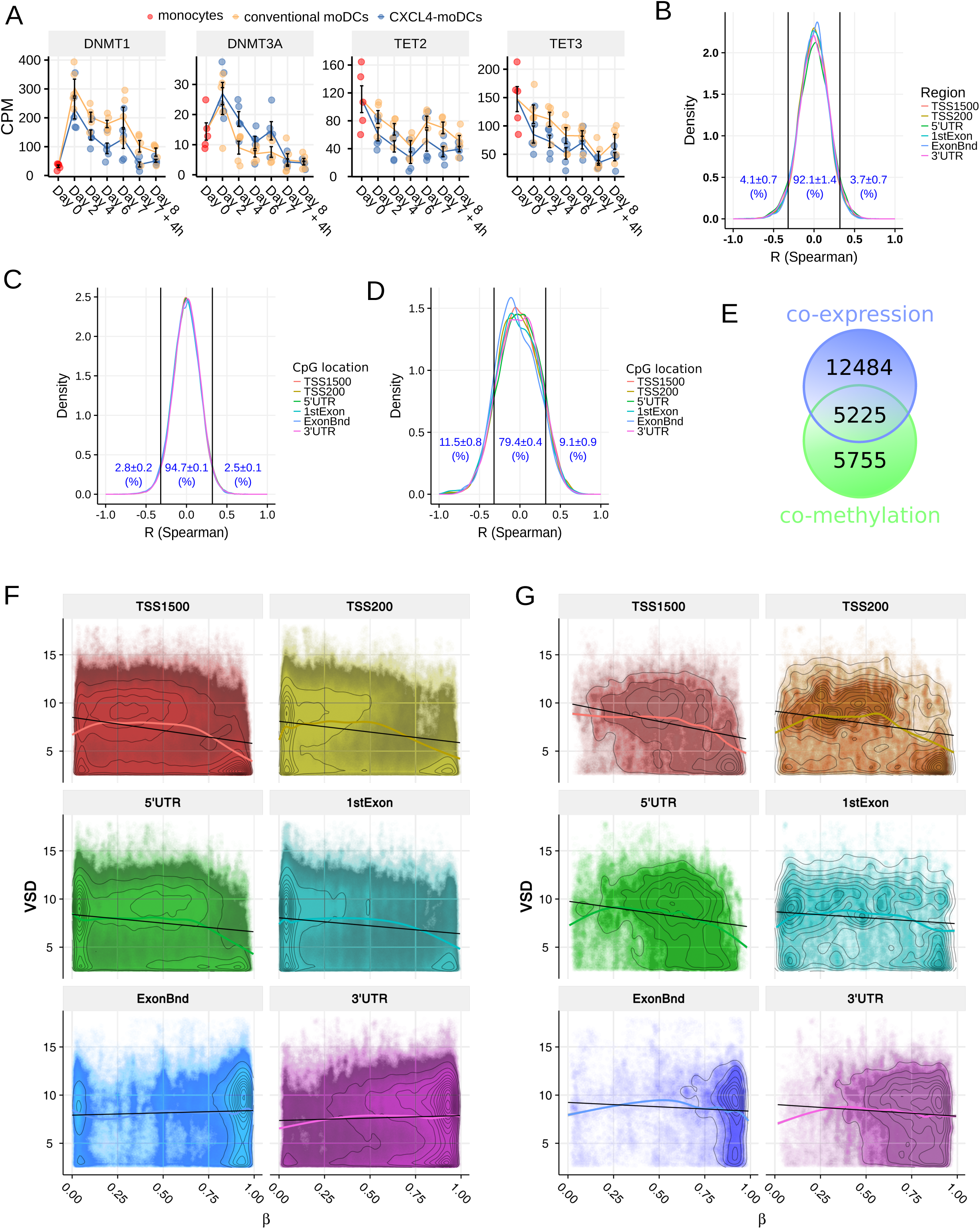
Comparison of transcriptome and DNA methylome. **(A)** Gene expression profile alterations of DNA methyltransferases and DNA demethylases. Data are shown as mean of count per million (CPM) ± SEM. **(B)** The distribution of Spearman correlation coefficients between the methylation levels of all regions (including the regions that are not differentially methylated) and the corresponding gene expression. **(C)** The distribution of Spearman correlation coefficients between the methylation levels of all CpG sites (including the CpGs that are not differentially methylated) and the corresponding gene expression. **(D)** The distribution of Spearman correlation coefficients between the methylation levels of differentially methylated CpGs and the corresponding gene expression. In (**B** to **D**) the cut-offs (two vertical lines at R = ±0.32) indicate significant correlation coefficients (*P* < 0.01). Overlap between differentially expressed genes (DEGs) and differentially methylated genes (DMGs) during differentiation and upon polyI:C stimulation. A gene is considered differentially methylated if there is at least one region within this gene that is differentially methylated. **(E)** Venn diagram shows the overlapping genes between DEGs and DMGs in all comparisons which were used for further analysis. **(F)** Contour plots show global comparison of β values (x-axis) and VSD values (y-axis) for all the genes. **(G)** Contour plots show global comparison of β values (x-axis) and VSD values (y-axis) for genes that are differentially expressed and methylated. The black straight lines were obtained by fitting a linear regression model and the smoothing curves were obtained by fitting a non-linear model (see Methods).

**Figure 3—figure supplement 2.**
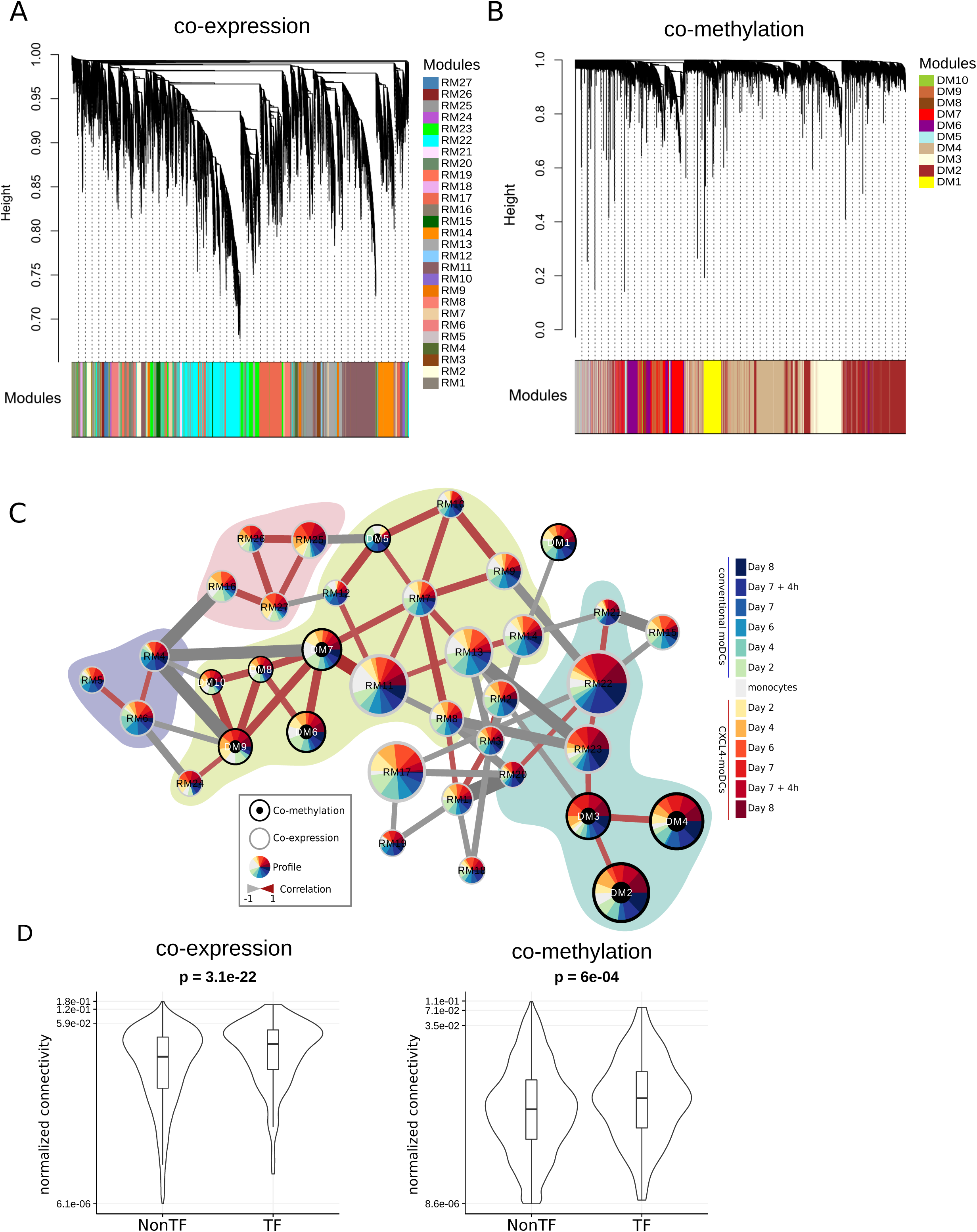
Modules of co-expression and co-methylation networks. Hierarchical clustering dendrogram of genes generated using topological overlap matrix (TOM) obtained from **(A)** gene expression or **(B)** methylation data. The co-expression and co-methylation modules were obtained using WGCNA package and are shown with different colors independently. **(C)** Network of co-expression and co-methylation modules based on the correlation of module eigengenes. Each circle (node) represents either co-expression or co-methylation module. Co-methylation modules are denoted as a node with a black circle in the middle, while the other nodes denote co-expression modules. The size of each node depicts the number of genes within that module. Different time and conditions are represented by the colors shown in the legend. The colored pie chart within each node represents the eigengene profile of that module. The edges (lines between two nodes) represent spearman correlation coefficient (ρ) of eigengenes between two modules. The thickness of edge depicts the absolute value of ρ and edges with absolute value of ρ<0.65 are not shown. Colored shades in the background depict strongly positively correlated modules. **(D)** Violin/box plots comparing the connectivity, normalized by the total connectivity in the corresponding module, for transcription (co-)factors (TF) and other genes (NonTF) in the co-expression (left) and co-methylation (right) networks.

**Figure 3—figure supplement 3.**
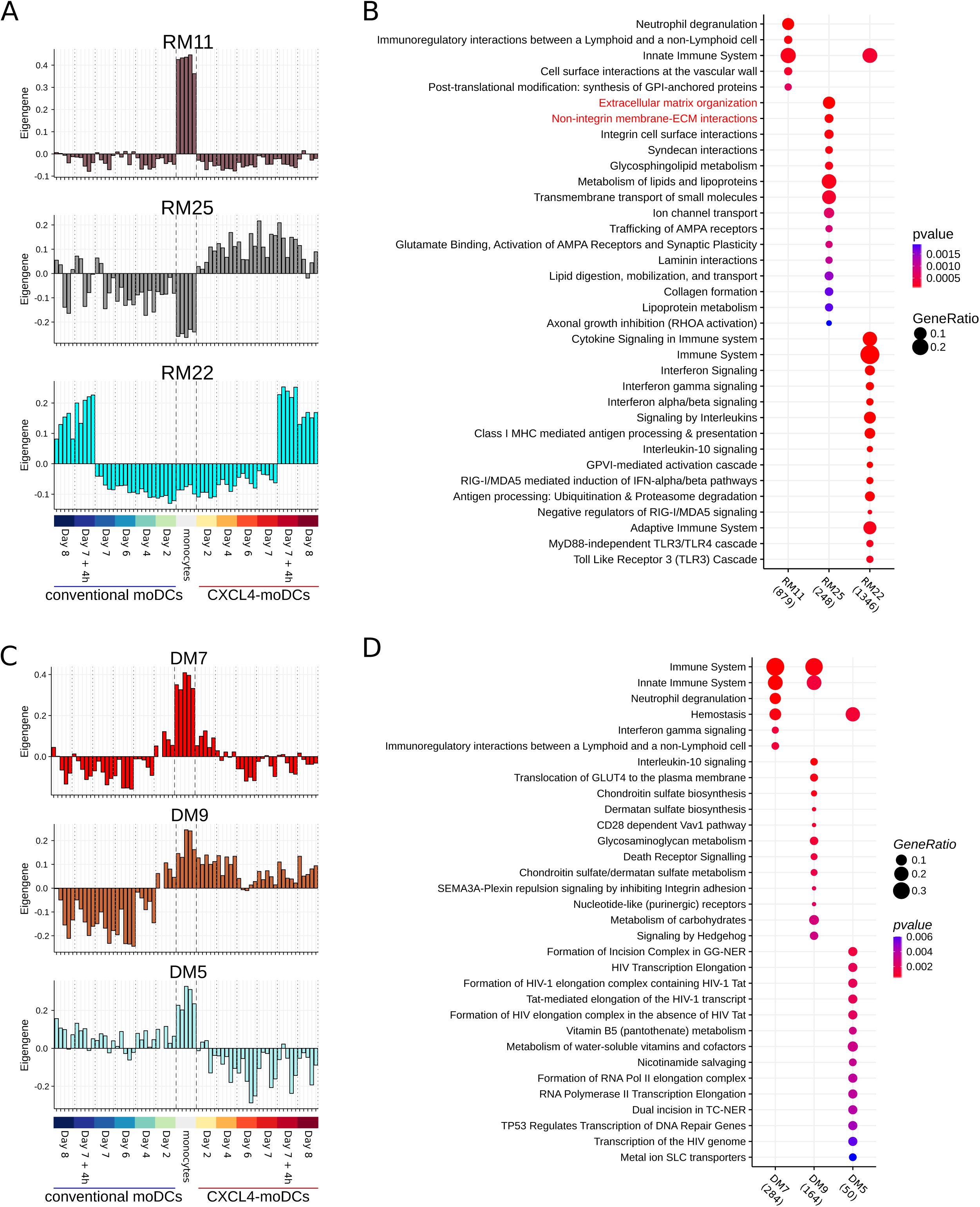
Characteristics of co-expression and co-methylation modules. **(A)** Bar charts show the eigengene of representative co-expression modules. **(B)** Pathway enrichment analyses for the modules shown in **(A)**. **(C)** Bar charts of the eigengene of representative co-mehtylation modules. **(D)** Enriched pathways analysis for the modules shown in **(C)**. In **(B)** and **(C)** the size of the circle depicts the gene ratio of DEGs in the pathway to the total number of DEGs in each set of genes; the colors of the circle represent FDR adjusted p values. In brackets, below the graph, we show the number of genes in each module. The full lists of pathways are available in **Supplementary File 8**.

**Figure 4—figure supplement 1.**
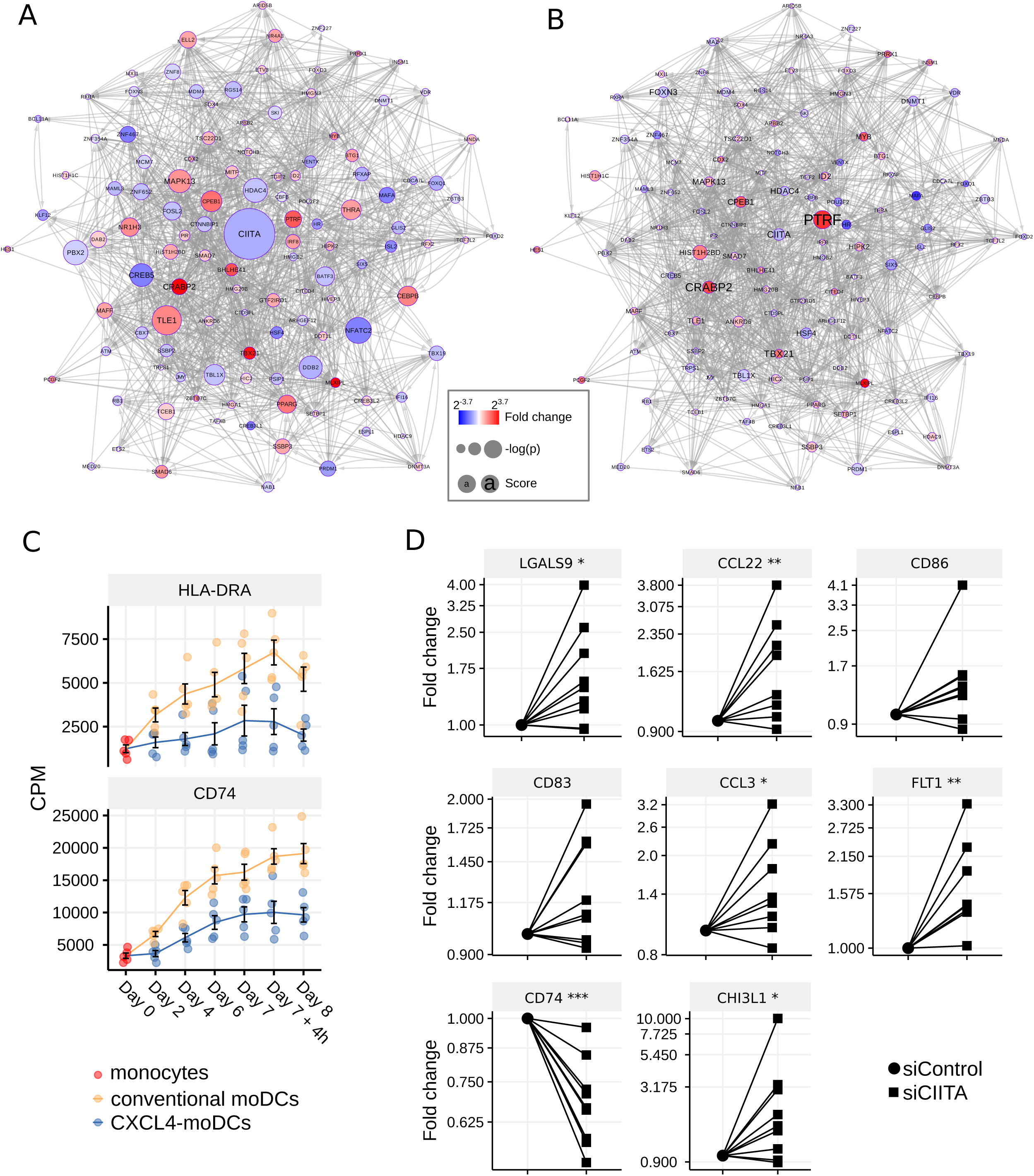
Enriched transcription regulators obtained using random forest based gene regulatory network. Enriched transcription regulator networks **(A)** during differentiation and **(B)** upon stimulation. Red indicates up-regulation and blue indicates down-regulation of TFs in CXCL4 moDCs compared to moDCs. Circle size shows (-log_10_(p)) obtained using differential expression analysis. Text size represents the overall score of TF enrichment obtained from RegEnrich (see Methods). Edges (lines) were obtained from top 5% edges in gene regulatory network inferred by random forest. To make the networks comparable same scales and parameters were used. **(C)** Gene expression profile of DEGs downstream to CIITA regulation. **(D)** After transfection of monocytes with Silencer negative control siRNA (siControl) and Silencer CIITA siRNA (siCIITA), cells were differentiated into moDCs for 6 days. On day 6, inflammatory and co-stimulatory genes were analyzed by qPCR. (Data are normalized by the mean expression of *RPL32* and *RPL13A*; fold change relative to siControl). Each symbol represents an individual moDC donor; lines connect the same donor. Paired two-sided Student’s *t*-test. * *P* <0.05, ** *P* < 0.01, *** *P* < 0.001.

**Figure 5—figure supplement 1.**
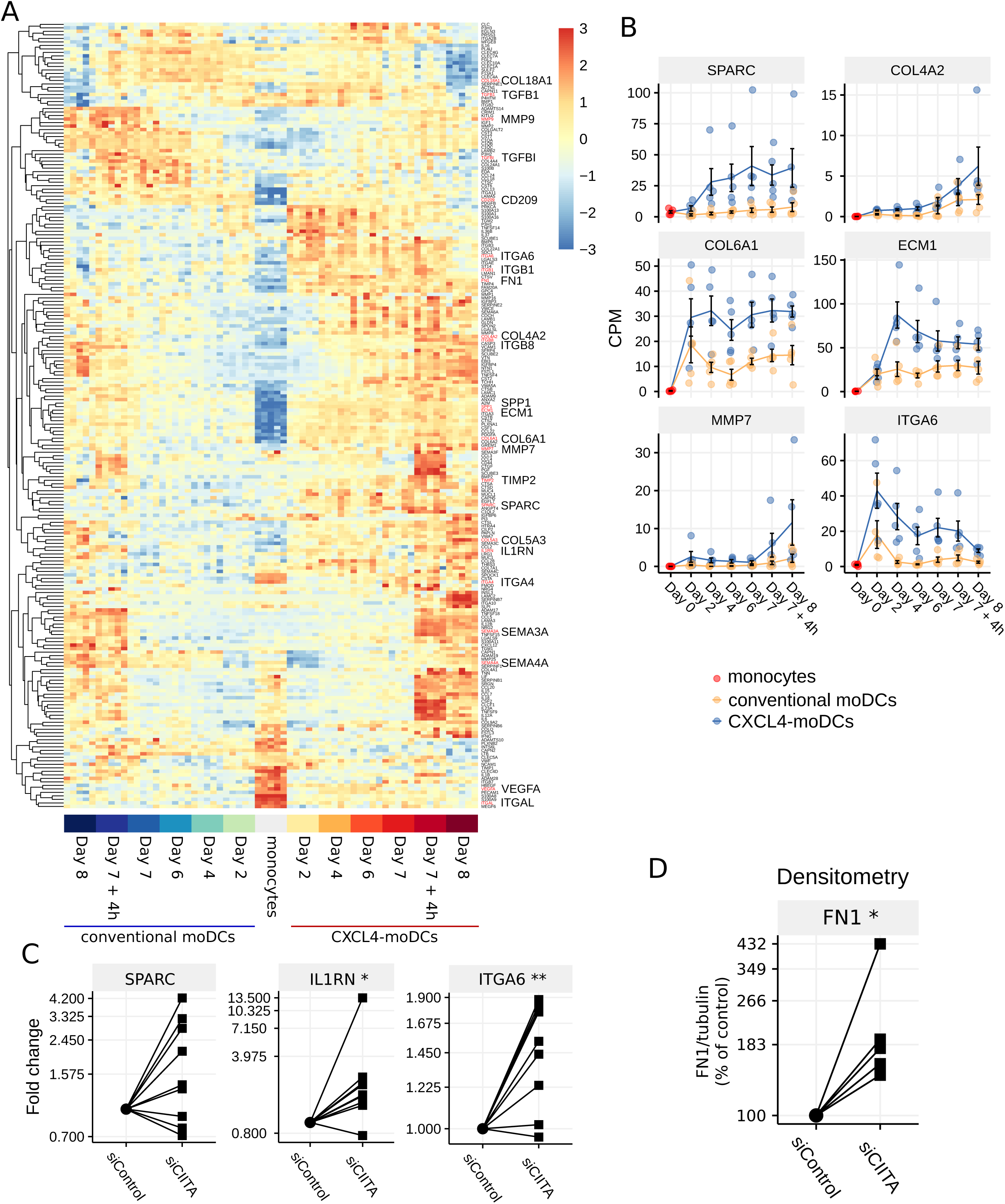
Identification of genes implicated in ECM remodeling. **(A)** Heatmap showing differentially expressed genes that play a role in ECM remodeling, identified from pathway enrichment analysis (**Figure 1—figure supplement 1B** and **Figure 1—figure supplement 2B**), and CXCL4 responsive co-expression modules (**Figure 3—figure supplement 2B**). Each column represents a sample and the colors on the bottom denote different time and conditions. The color schemes in the heatmap are shown as z-scores. **(B)** Expression profiles of example genes implicated in ECM remodeling. Data are shown as mean of count per million (CPM) ± SEM. **(C)** After transfection of monocytes with Silencer negative control siRNA (siControl) and Silencer CIITA siRNA (siCIITA), cells were differentiated into moDCs for 6 days. Genes involved in ECM remodeling/fibrosis analyzed by qPCR on day 6. Data are normalized by the mean expression of RPL32 and RPL13A; fold change relative to siControl. **(D)** Fibronectin (FN1) and tubulin expression measured by western blot on day 6. Signal intensity of 5 independent experiments was quantified by densitometry analysis. To determine the relative expression of FN1 between siControl and siCIITA, the ratio between the expression of FN1 and tubulin was first calculated. In panels **(C)** and **(D)** each symbol represents an individual moDC donor; lines connect the same donor. Paired two-sided Student’s *t*-test. * *P* <0.05, ** *P* < 0.01, *** *P* < 0.001.

**Figure 5—figure supplement 2.**
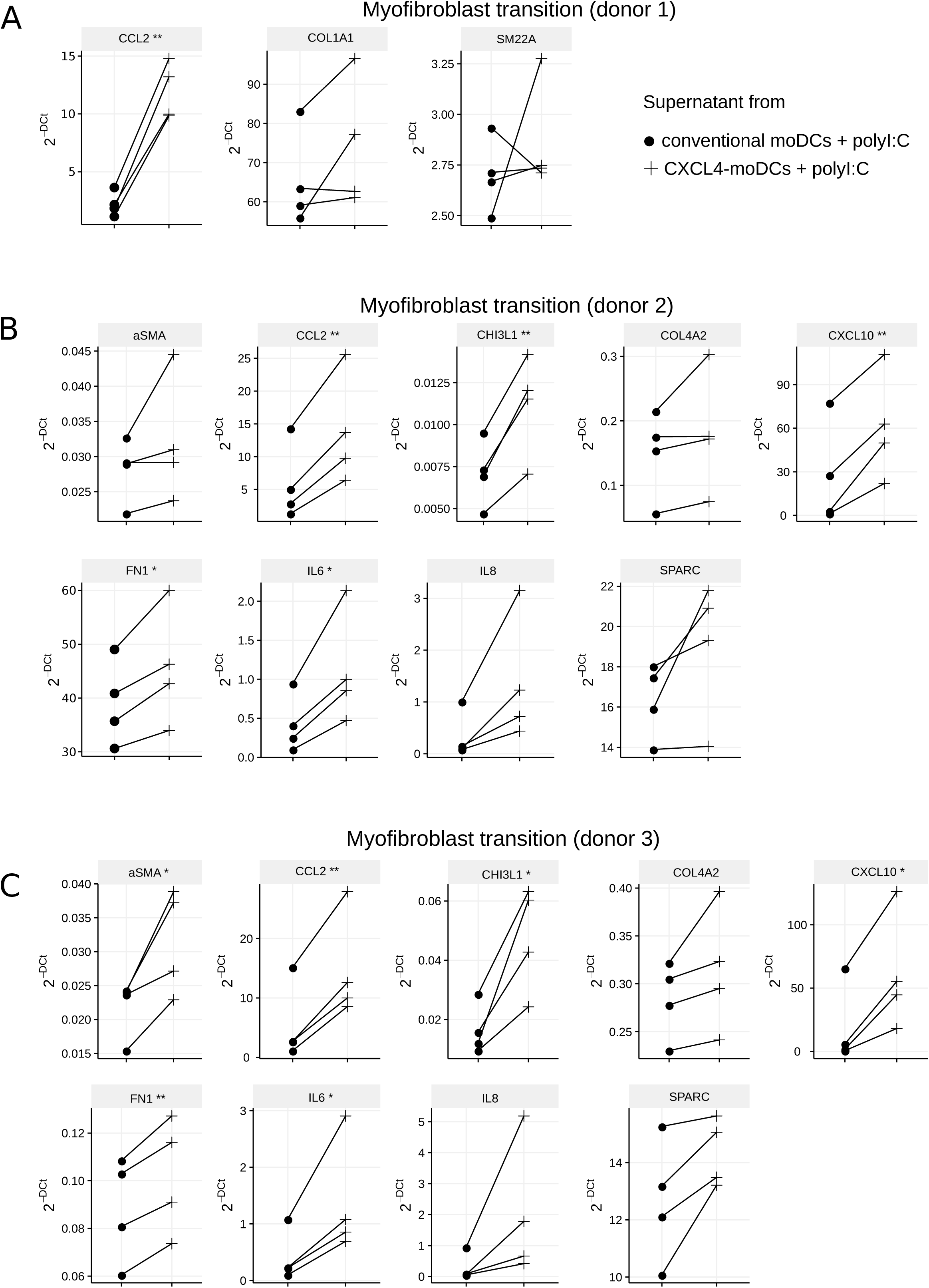
Gene expression analysis of healthy myofibroblasts after exposure to cell-free supernatants from CXCL4 moDCs and moDCs. CXCL4 moDCs and moDCs on day 7 were stimulated with polyI:C for 24 hours. Cell-free supernatants were added to healthy myofibroblasts for 24 hours. Inflammatory and fibrotic gene expression was analyzed by qPCR. Each symbol represents an individual moDC donor (n=4). Lines connect the same moDC donors. **(A)** shows the gene expression for the first fibroblast donor. The remaining measured genes are shown in Figure 5H. The gene expression for the **(B)** second and **(C)** third independent fibroblast donors. Paired two-sided Student’s *t*-test. * P <0.05, ** P < 0.01, *** P < 0.001.

## Supplementary Files

**Supplementary File 1.**

The number of differently expressed genes (DEGs) and differently methylated genes (DMGs) during differentiation and upon polyI:C stimulation.

**Supplementary File 2.**

Primer list for RT-qPCR.

**Supplementary File 3.**

DEGs and p-values by differential expression analysis.

**Supplementary File 4.**

DMRs and p-values by differential methylation analysis.

**Supplementary File 5.**

Full list of enriched pathways for the DEGs during differentiation.

**Supplementary File 6.**

Full list of enriched pathways for the DEGs upon stimulation.

**Supplementary File 7.**

Full list of enriched pathways for the DEGs overlapping during differentiation and upon stimulation.

**Supplementary File 8.**

Full list of enriched pathways for co-expression and co-methylation modules.

**Supplementary File 9.**

Transcription regulators used as TFs for the analysis.

